# Ecological niche differentiation mediates near complete premating reproductive isolation within the *Gladiolus carneus* (Iridaceae) species complex

**DOI:** 10.1101/2025.04.04.647161

**Authors:** Katharine L. Khoury, Shelley Edwards, Ethan L. Newman

## Abstract

**Background and aims:** Ecological niche differentiation is well associated with intraspecific divergence of functional traits, which may lead to the evolution of premating reproductive isolation. However, the link between the ecological niches, trait divergence, and premating isolation remains poorly understood. This is particularly pertinent in hyper-diverse areas such as the Cape Floristic Region (CFR) of South Africa, where fine-scale ecological heterogeneity has been hypothesised as a major driver of speciation. Using the polymorphic geophyte *Gladiolus carneus,* endemic to the CFR, we test whether ecological niche differentiation mediates premating reproductive isolation.

**Methods:** We first tested whether newly and previously described *Gladiolus carneus* varieties were distinct based on their floral and vegetative morphology. Next, we documented each variety’s abiotic niche, flowering phenology and pollination niche and further tested whether any resulting niche differentiation causes premating reproductive isolation.

**Key results:** Seven morphologically distinct varieties were identified, of which two of these are newly documented. Using niche modelling and multivariate analyses, we found that these varieties occupied distinct abiotic niches, resulting in strong ecogeographic isolation. They also had distinct flowering times, causing varying strengths of phenological isolation. For the pollinator niche, we found that all sampled populations had a single highly effective functional pollinator; however, at the variety level, there were no consistent trends leading to varying strengths in pollinator-mediated isolation. Across all varieties, ecogeographic isolation was the strongest gene flow barrier, which combined with phenological and pollinator-mediated isolation, causes near complete premating reproductive isolation.

**Conclusions:** These results suggest that ecological niche differentiation between *Gladiolus carneus* varieties may be contributing to incipient speciation within the species complex and further suggests that ecological niche differentiation may be a major driver of speciation in the hyper-diverse Cape Floristic Region.

## INTRODUCTION

At the early stages of ecological speciation, ecologically-based divergent selection is key in producing phenotypes that often associate with distinct biotic and abiotic niches (Schluter, 2000, Schluter, 2001, Rundle and Nosil, 2005). These divergent niches, which distinct phenotypes occupy, are often directly associated with gene flow barriers that act before mating (Coyne and Orr, 2004, Rundle and Nosil, 2005). In seed plants, these premating barriers are thought to be important in completing reproductive isolation, especially in recently diverged taxa or ecotypes (Christie *et al*., 2022), making it pertinent to disentangle the ecological shifts underlying barrier effectiveness, when studying lineage diversification. For example, if shifts in abiotic factors cause divergent selection on the functional traits of a plant in different parts of its range, it will likely result in ecogeographic isolation, which is defined as a reduction in encounter rates due to spatial separation as a result of genetically based differences (Sobel, 2014). Similarly, closely related taxa associated with divergent pollinator niches may have divergent floral phenotypes that promote pollinator-mediated isolation through unique pollinator preferences and morphological fit with flowers (Kay, 2006, Sobel and Streisfeld, 2015, Minnaar *et al*., 2019). Other phenotypic shifts, such as flowering times, are difficult to associate with any single biotic or abiotic driver. Shifts in flowering phenology can be due to either selection by biotic and abiotic factors (Elzinga *et al*., 2007, Munguia-Rosas *et al*., 2011), or by plastic responses to the abiotic environment (Elzinga *et al*., 2007). Regardless of the causes of the shift in flowering times, it can result in phenological isolation (Paudel *et al*., 2018, Ramirez-Aguirre *et al*., 2019), which is a reduction in gene flow due to a mismatch in flowering times (Coyne and Orr, 2004).

Although, ecological shifts in association with early lineage diversification have been well documented (Grossenbacher *et al*., 2014, Dellinger *et al*., 2024, Fernandez-Mazuecos and Glover, 2024), this is rarely placed in a reproductive isolation context (*but see* Sandstedt *et al*., 2021, Boucher *et al*., 2023, Farnitano and Sweigart, 2023). Usually, reproductive isolation is studied by either quantifying the strength of multiple barriers between sister taxa (*see* Ramsey *et al*., 2003, Kay, 2006, Sobel and Streisfeld, 2015, Ostevik *et al*., 2016, Christie and Strauss, 2019, Ivey *et al*., 2023) or by focussing on a single barrier across multiple taxa pairs (Sobel, 2014). The first approach provides valuable information on the relative strength of barriers between taxa pairs, while the second approach shows the role of single barriers in driving wider trends in reproductive isolation within and between lineages. If these approaches were combined, where multiple gene flow barriers were quantified to test their relative importance across multiple taxa pairs, it would show which barriers played the largest role in maintaining reproductive isolation and driving the diversification of lineages as a whole. This approach has highlighted the role of multiple postpollination barriers (particularly hybrid seed inviability) in reducing gene flow between *Mimulus* species in the section *Eunanus* (Farnitano and Sweigart, 2023), for postzygotic barriers completing reproductive isolation in the *Mimulus tilingii* species complex (Sandstedt *et al*., 2021), and for showing that geographic and phenological isolation, maintain species boundaries within the genus *Argyroderma* (Boucher *et al*., 2023).

Documenting ecological niche differentiation in association with premating isolation is particularly pertinent in hyper-diverse areas such as the Cape Floristic Region (CFR), where interactions between topography, soils, climate, and pollinators have generated ecological heterogeneity that has facilitated the diversification of plant lineages (Ellis *et al*., 2014). Specifically, the CFR is characterised by folded mountain belts that run parallel to the Indian and Atlantic ocean coasts (Linder, 2003), which gave rise to some of the most nutrient-poor soils currently recorded on Earth (Cramer *et al*., 2014). However, these soils contrast with the moderately fertile, shale-derived soils in coastal planes and intermontane valleys (Cramer *et al*., 2014), which have created a mosaic of edaphic conditions in the CFR. Rainfall varies along both an east-west and elevational gradient. In the west, there is a strongly seasonal Mediterranean climate, while in the east, the climate is much more aseasonal with rainfall throughout the year (Manning and Goldblatt, 2012). Further variation is created by steep mountainous landscapes, resulting in orographic rainfall (Manning and Goldblatt, 2012). These interacting conditions have created a heterogenous abiotic selective landscape that has facilitated frequent ecological speciation in the CFR (van der Niet and Johnson, 2009, Schnitzler *et al*., 2011, Ellis *et al*., 2014). Apart from diverse abiotic niches that have contributed to lineage diversification, shifts in biotic niches associated with pollinators (van der Niet and Johnson, 2009, Valente *et al*., 2012) are also thought to be important drivers of lineage diversification in the Cape. These ecological shifts considering both biotic and abiotic niches have been shown at a macroevolutionary scale, focusing on species-level phylogenies of the major Cape clades (van der Niet and Johnson, 2009, Schnitzler *et al*., 2011, Valente *et al*., 2012), as well as through trait-by-environment associations (Carlson *et al*., 2011, Mitchell *et al*., 2015, Newman *et al*., 2015). However, few of these ecological shifts have been directly linked to the strength of corresponding premating isolation barriers in the CFR (*but see* Newman and Johnson, 2024) to elucidate the relative importance of individual gene flow barriers to lineage diversification.

*Gladiolus carneus* Delaroche (Iridaceae) is a geophyte endemic to the CFR (Figure 1). The species complex is polymorphic and is made up of at least seven distinct ‘varieties’ that differ in both their morphological traits and geographic distribution (Lewis *et al*., 1972, Delpierre and du Plessis, 1974). Although, recognized by Delpierre and du Plessis (1974), the varieties are not well defined based on their functional traits. Specifically, both vegetative and floral traits, and particularly nectar guide properties, vary substantially between varieties. In addition to the varieties varying in functional traits, they also seem to occupy distinct ecological niches within discrete geographic ranges. This provides an opportunity to explore the relationship between trait divergence, ecological niche differentiation, and premating reproductive isolation in a diverging species complex within the CFR. In this study system, we first test whether previously described and two undescribed varieties are morphologically distinct from one another. Next, we ask do these varieties occupy distinct biotic and abiotic niches? and, are the differences in these ecological niches resulting in premating reproductive isolation between varieties?

**Figure 1.**
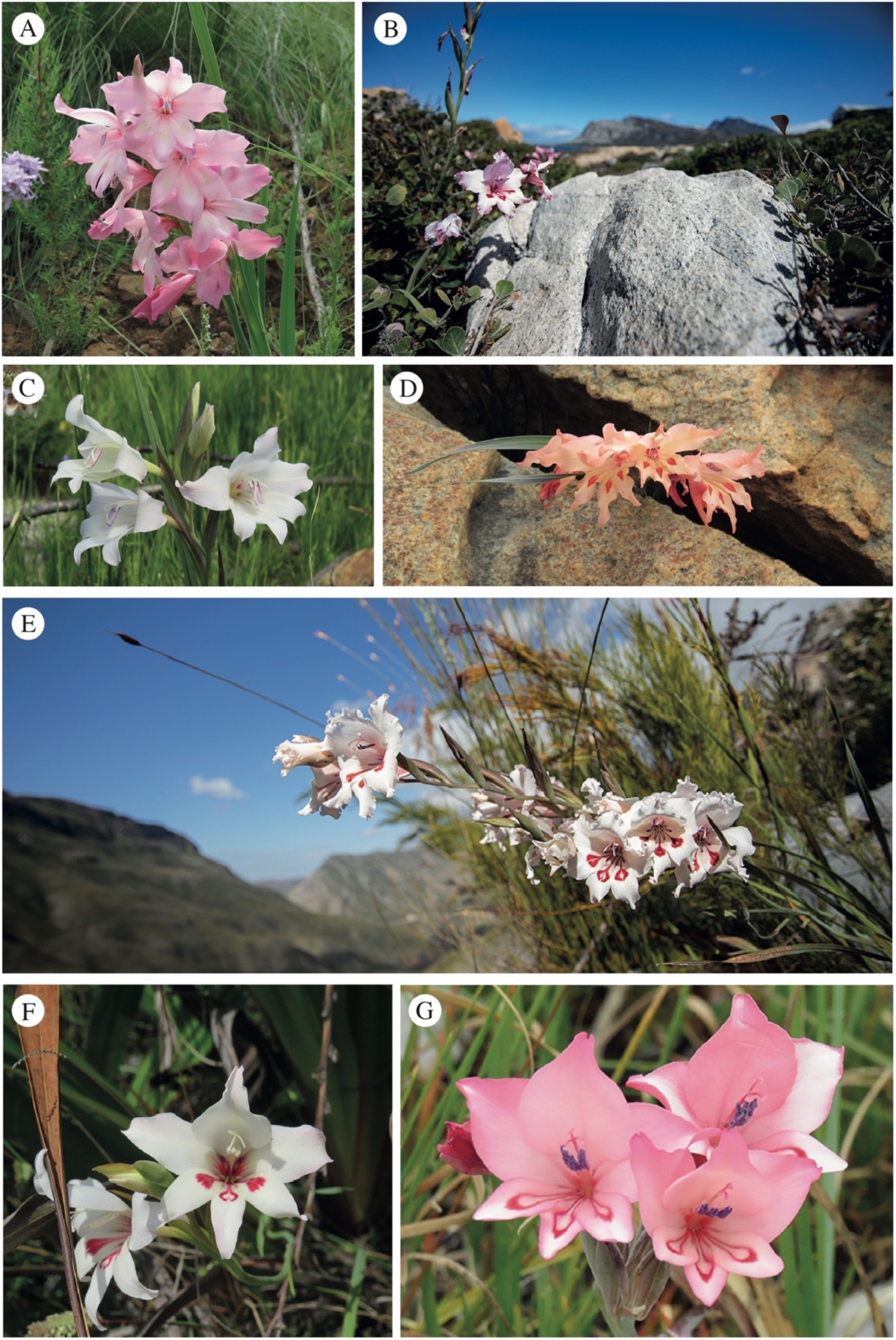
Colour plate of all *G. carneus* varieties. The varieties are (A) *albidus*, (B) *blandus*, (C) *callistus*, (D) *high-altitude*, (E) *langeberg*, (F) *macowanianus*, and (G) *prismatosiphon*. Photos: A, C, D, F, G: Katharine Khoury; B, E: Ethan Newman.

## MATERIALS AND METHODS

### Study species

*Gladiolus carneus* is a deciduous cormous geophyte endemic to the CFR. The species occurs from Ceres, south into the Cape peninsula, and east along the Southern Cape coast and then inland into the Langeberg and Outeniqua Mountains (Goldblatt and Manning, 1998). *Gladiolus carneus* occurs along the entire altitudinal gradient of the CFR, with populations occurring from sea level along the rocky Kleinmond coast, up to the high peaks of the South-Western and Southern Cape over 1000 meters above sea level. The species flowers from August into early January (Lewis *et al*., 1972). The species has been described under at least 20 different names, with many of those taxonomic distinctions referring to specific populations or forms of the species. These forms were eventually sunk into the single species, *G. carneus* by Lewis *et al*. (1972). However, the species can largely be defined into a number of distinct ‘varieties’ based on their morphology and geographic distribution (Delpierre and du Plessis, 1974). Within this study, the varieties include *albidus, blandus, callistus, macowanianus,* and *prismatosiphon* as defined by morphology and localities described by Delpierre and du Plessis (1974). The form of *G. carneus* that has been found in the Langeberg mountains has been grouped in both *blandus* (Delpierre and du Plessis, 1974) and *prismatosiphon* (Lewis *et al*., 1972). However, due to its distinct geographic range and floral morphology, it has been treated as a separate variety within this study. The late-flowering, high-altitude forms of *G. carneus* in the Drakenstein, Hottentots-Holland and Riviersonderend mountain ranges were treated as a distinct *high-altitude* variety. *Albidus* (Figure 1A) occurs in Paarl, Stellenbosch, and down into Hermanus (Delpierre and du Plessis, 1974). The variety can be identified by the white flowers with pale yellow markings on the lower tepals (Delpierre and du Plessis, 1974). *Blandus* (Figure 1B) is found near Kleinmond (Delpierre and du Plessis, 1974). *Callistus* (Figure 1C) has unmarked flowers with a deep purple gullet (Delpierre and du Plessis, 1974). *Macowanianus* (Figure 1F) is the most typical form of the species, found in the Southern Cape and along the Cape peninsula (Delpierre and du Plessis, 1974), and identified by distinct dark nectar guides. *Prismatosiphon* (Figure 1G) is a form found solely on rocky outcrops near Bredasdorp (Delpierre and du Plessis, 1974). There is also a form of *G. carneus* that has been found in the Langeberg and Outeniqua mountains (Figure 1E). The *high-altitude* variety includes all late-flowering populations occurring on the top of the Drakenstein, Hottentots-Holland, and Riviersonderend mountain ranges (Figure 1D), which are frequently exposed to freezing temperatures in the winter.

### Are previously and newly described varieties of *Gladiolus carneus* morphologically distinct?

#### Morphological differences between varieties of Gladiolus carneus

To test whether *G. carneus* separates into distinct varieties based on morphology, we measured morphological traits in 28 populations (*see Table S1 for site coordinates*) representing the entirety of the geographic and morphological variation within the species complex (*see Table S2 for sample sizes*). This includes populations that are representative of all previously described varieties [*sensu* Delpierre and du Plessis (1974)], and two the newly identified varieties. On each individual, some or all of the following morphological traits were measured using digital callipers (0-200mm, TA): floral tube length, flower gape, petal size, flower width, and width of the longest leaf, while inflorescence height and the length of the longest leaf was measured using a measuring tape (Figure S1A-C). We also documented the total number of flowers and leaves on each individual. Flower gape was measured from the lowest anther to the lower median tepal directly underneath (Figure S1A). Flower width was the distance between the tips of the lateral tepals (Figure S1A). Floral tube length was measured from the top of the ovary to the notch connecting the corolla to the lower tepals (Figure S1B). Petal size was measured from the tip to the base of the dorsal tepal (Figure S1B). Inflorescence height was measured from the base of the inflorescence to the top the display (Figure S1C). Leaf length was measured as the distance from the ground to the tip of the longest leaf (Figure S1C), whereas leaf width was measured as the widest point on the longest leaf (Figure S1C). The total flowers included all buds, open and senesced flowers, and the total number of leaves included all leaves at the base of the fan.

All data analysis was conducted in R v.4.1.2 (R Core Team, 2021). We used a series of Principal Component Analyses (PCA’s) to determine if different varieties cluster separately based on their vegetative and floral morphology. We implemented the PCAs using the *prcomp* function from the ‘*stats*’ package (R Core Team, 2021) and tested whether the varieties cluster separately using a permutation MANOVA with 999 permutations. We then used a pairwise permutation MANOVA with a Bonferroni correction for pairwise comparisons between varieties. The permutation MANOVA was implemented using the *adonis2* function in the package ‘*vegan*’ (Oksanen *et al*., 2020) and we used the *pairwise.perm.manova* from the ‘*RVAideMemoire*’ package (Herve, 2023) for pairwise comparisons. We used this procedure to test whether the varieties separate based on their floral traits, vegetative traits, and all morphological traits.

We further determined if there were significant differences between varieties using generalized linear models (GLMs) implemented in the ‘*stats*’ package (R Core Team, 2021) for the following morphological traits: tube length, flower gape, petal size, inflorescence height, length and width of the longest leaf. These traits were chosen as they represent functional traits directly linked to the biological variation associated with their biotic and abiotic niches (e.g., tube length is associated with pollinator proboscis length). In the models, each trait was set as the response variable while the explanatory variables were variety, and site nested within variety. The nested term was included to account for sampling at multiple sites per variety. All traits were fitted with either a Gaussian or Gamma error distribution, depending on the normality of the dataset. Significance of model terms were obtained using the *Anova* function from the ‘*car*’ package (Fox and Weisberg, 2019) while the *emmeans* function from the ‘*emmeans*’ package (Lenth, 2022) was used for pairwise comparisons.

#### Colour differences between varieties of Gladiolus carneus

To test if there were differences in the colour traits between varieties of *G. carneus* within bee and fly colour vision, we sampled 1266 spectral measurements from 300 individuals across 24 populations (*see Table S1 for site coordinates*) in 2020, 2022, and 2023 (*see sample sizes in Table S3*). For each individual of *macowanianus*, *blandus, prismatosiphon, high-altitude* and *langeberg* varieties, we measured the spectral reflectance of the dorsal tepal, the outline and centre of the nectar guides (hereafter referred to as the ‘guide’ and ‘centre’) on the lower median tepal and lower lateral tepal (Figure S1D). Only the dorsal tepal and centre of the nectar guides were documented for *albidus* (Figure S1E), as the variety lacks well-developed outlines around the nectar guides. As *callistus* lacks any nectar guides, only the dorsal tepal and the gullet for each individual was measured (Figure S1F). All spectra collected in 2020 was measured using an Ocean Optics S2000+ spectrometer with a DT-mini light source and fibre optic probe (UV/VIS 400 μm), while spectra collected in 2022 and 2023 were measured using Ocean Insight FLAME Miniature spectrometer (Ostfildern, Germany) with a PX-2 Pulsed Xenon Light Source and a Premium 400 μm fibre optic probe. All varieties’ spectra were processed, and negative values were corrected using the *procspec* function for each variety. All problematic spectra were removed. Aggregation plots showing the mean and standard error of the varieties for each trait were generated using the *aggplot* function to visualise the curvature of the reflectance spectra for each species (Figure S2).

Colour vision models were used to determine how pollinators perceive the colours of each variety. Reflectance spectra of each variety were plotted in the trichromatic colour hexagon (Chittka, 1992) for the *Apis mellifera* visual system and the colour-opponent coding vision model for flies (Troje, 1993). Bees have a conserved trichromatic visual system with UV, blue and green photoreceptors (Briscoe and Chittka, 2001), which are represented as vertices in the Chittka (1992) trichromatic colour hexagon. When spectra are plotted into the colour hexagon, spectra that fall into different quadrants are considered distinguishable by bees (Chittka, 1992). However, bees also have continuous colour discrimination, and they can reliably detect differences between colours within the same quadrant for distances larger than 0.11 hexagon units (Dyer, 2006), although with absolute conditioning, they can start to detect distances at 0.06 hexagon units (Dyer, 2006).

Fly colour vision was modelled using the colour-opponent coding (COC) model (Troje, 1993). The COC model is based on behavioural experiments with *Lucilia* blowflies that showed their colour vision was based on an opponency mechanism involving pairs of photoreceptors, namely R7p with R8p and R7y and R8y. *Lucilia* flies distinguish colour categories that depend on the relative excitation of the paired receptors. The COC model uses the relative quantum catches of the four fly photoreceptors to plot spectra into quadrants in a Cartesian plane. Loci plotted within a quadrant are too similar to be perceived, while loci in different quadrants are considered perceptible colour differences. However, Hannah *et al*. (2019) found that the hoverfly, *Eristalis tenax*, can discriminate between colours within the COC quadrants, and based on these experiments, Garcia *et al*. (2022) proposed four behaviourally relevant colour discrimination thresholds for *E. tenax*. From their study, we used 0.096 units, the lower boundary of ‘easily discriminable’ and 0.021, the lower boundary of ‘functionally discriminable’, to measure colour discrimination (Garcia *et al*., 2022). As the spectral sensitivities of fly pollinators of *G. carneus* are currently unknown, we used the spectral sensitivities of *E. tenax*, a widespread nectar-feeding fly.

Using the spectra from each part of the flower, quantum catches for both the bee and fly models were calculated with the following formula:

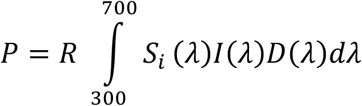

where *S_i_* is the sensitivity of each photoreceptor, *I*(*λ*) is the spectral reflectance and *D*(*λ*) is the daylight illuminant. Quantum catches were hyperbolically transformed using:

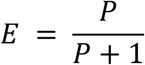

and plotted into hexagonal space using:

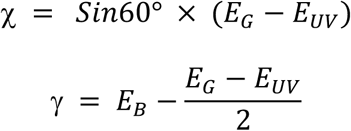

to find the x and y coordinates. While the formulas:

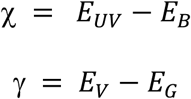

were used to find the coordinates of the relative quantum catches in categorical fly space. All visual modelling used D65 daylight illuminant and green foliage background.

We used a MANOVA on the cartesian coordinates within the colour vision models to test for statistical differences between the varieties for each colour trait. This approach accounts for the multivariate nature of the data (Maia and White, 2018). We used the *mvpaircomp* function with a bonferroni correction from the *‘biotools’* package (da Silva *et al*., 2017) for pairwise comparisons between the varieties for each colour trait. In cases where there were statistical differences between the varieties within bee and fly colour vision, We then tested if the differences were perceptible (Maia and White, 2018). Empirical means and bootstrapped confidence intervals of colour distances between varieties were generated using the *bootcoldis* function (Maia and White, 2018). The differences between varieties were considered to be distinguishable if the mean was above the discrimination threshold of 0.11 hexagon units in bee colour vision and above 0.096 Troje units within fly colour vision. All spectral processing, visualisation and colour vision modelling were completed using the R package ‘*pavo’* (Maia *et al*., 2019)

### Do *Gladiolus carneus* varieties occupy distinct biotic and abiotic niches?

#### Point locality sampling from iNaturalist

We used iNaturalist observations to extract flowering and point locality data for all *G. carneus* varieties. The flowering data was used to test for differences in the phenological niche between varieties, and the point localities were used to extract abiotic data from environmental layers that test for differences in the realized and fundamental abiotic niche between varieties. We used iNaturalist observations as *G. carneus* is well documented (1500+ observations as of September 2024). The observations include images of the flowers allowing for reliable identification of the varieties, and the associated locality and accuracy data provides reliable occurrence data across its geographic range. Additionally, *G. carneus* is documented yearly on iNaturalist allowing for multi-year observation dates that can be used to capture temporal variation in flowering times between years and localities. We classified iNaturalist observations into the seven varieties based on their floral morphology and location. Furthermore, we excluded any observations that (1) did not have clear photos of the flowers, (2) observations of closely related taxa that had been misidentified as *G. carneus,* (3) observations that did not fit the descriptions of previously described varieties or were likely hybrids, (4) observations that occurred outside of the natural range of the species within the Cape Floristic Region (e.g., Australia), and (5) observations that were not research grade.

#### Do varieties of Gladiolus carneus occupy distinct abiotic niches?

To test whether the varieties of *G. carneus* occupy distinct abiotic niches, we modelled the predicted distribution of each variety using MaxEnt (Phillips *et al*., 2006, Elith *et al*., 2010). All iNaturalist observations were filtered to include coordinates with an open geoprivacy and an accuracy under 100 m to ensure that the niches of each variety were accurately represented at fine spatial scales. In addition to iNaturalist data, we provided point localities for varieties that were not well documented from observations in the field (K. Khoury, unpublished data). Overall, we documented 209 *albidus*, 33 *blandus*, 67 *callistus*, 34 *high-altitude*, 37 *langeberg*, 379 *macowanianus* and 27 *prismatosiphon* point localities to be used in subsequent analyses. We mined 19 bioclimatic layers and an elevation layer at the 30 arc second (∼1km) resolution from WorldClim (https://www.worldclim.org/data/worldclim21.html), while seven soil layers were taken from Cramer *et al*. (2019). The soil layers from Cramer *et al*. (2019) were modelled specifically in the Cape Floristic Region and allow for more reliable predictions of vegetation type in the CFR than the comparable SoilGrids layers. All stacking, resampling, and processing of abiotic layers was done using the ‘*raster*’ package (Hijmans, 2024). The *res* function was used to ensure resolution of all WorldClim and soil variables matched. All abiotic layers were resampled to match the distribution of the soil layers using the *resample* function. The resampled abiotic layers covered the entire native range of *G. carneus*. The collinearity of all abiotic variables was tested using a Pearson correlation and any abiotic variables with a correlation coefficient >|0.69| were excluded from further analysis. In total, we were left with nine biologically relevant, uncorrelated abiotic variables that were used in the final models (*see Table S4 for descriptions of all layers*). We constructed maximum entropy species distribution models for each variety using the *maxent* function in the package ‘*dismo’* (Hijmans *et al*., 2023). The models was implemented using 75% of the occurrence data for model training and 25% was used for model testing, with 10 000 background points (Elith *et al*., 2010). The *evaluate* function was used to calculate AUC scores to assess model performance. AUC scores range from 0 to 1, with scores closer to 1 indicating better model performance. The equal sensitivity and specificity threshold was used to create binary maps from the probability predictions of the MaxEnt output using the *threshold* and *reclassify* functions. Values above the threshold were counted as present, while values below were counted as absent. A jackknife test was used to determine the most important environmental variable in each model.

We extracted abiotic layers for each point locality to test whether the varieties occupied distinct realised abiotic niches. We used a PCA to test for niche differentiation, which included the uncorrelated variables used in the niche modelling. We used a permutation MANOVA to test if the varieties cluster separately based on their abiotic niche and then a pairwise permutation MANOVA with a Bonferroni correction for pairwise comparisons between the varieties. The PCA, permutation MANOVA and pairwise permutation MANOVA was implemented using the same procedure described under the morphological analysis.

#### Do varieties of Gladiolus carneus flower at different times?

iNaturalist observations were used to document differences in flowering times across all varieties. As the total number of flowers are often not documented on iNaturalist observations, the total observations per day was used as a proxy for the flowering times. In total, there were 307 *albidus*, 59 *blandus*, 129 *callistus*, 51 *high-altitude*, 38 *langeberg*, 612 *macowanianus*, and 33 *prismatosiphon* observations used in the temporal analysis.

As flowering data is a seasonal event, best represented as circular data, we used circular statistics to test for differences in flowering times between the *G. carneus* varieties (Pewsey *et al*., 2013). To facilitate the analysis, each day of the year (1 to 365) was converted into radians, the standard unit of angular measurement. The number of observations against each radian was then used in the analysis. A Mardia-Watson-Wheeler test, a non-parametric test of the homogeneity of two or more samples of circular data, was used to test for significant differences in the flowering times between varieties. We used a series of Mardia-Watson-Wheeler tests to conduct pairwise comparisons between the varieties’ flowering times. A Bonferroni correction was applied to the resulting p-values to reduce the likelihood of Type I errors. All analysis was conducted using the *watson.wheeler.test* function in the ‘*circular’* package (Agostinelli and Lund, 2024).

#### Do varieties of Gladiolus carneus occupy distinct pollination niches?

Pollinator observations were conducted across 16 sites of *G. carneus* in 2020, 2022 and 2023 to identify the main pollinators of each variety. A total of 723 flowers were observed over more than 69 hours (*see breakdown of observations per site in Table S5*). Observations were conducted between 8.30am and 2.30pm on days above 20° Celsius. At each site, a maximum of two observers recorded all pollinator visits, the total observed flowers, and the total observation time. All visits recorded on focal plants within the observation time were used to the calculate visitation rates (visits.flower^-1^.hour^-1^) of pollinators at each population. In addition to recording visits, pollinators were caught using an insect net and euthanized using ethyl acetate fumes. These insects were used for species level identification, the measurements of pollinator functional traits, as well as counting conspecific pollen loads. On each insect we measured extended proboscis length and thorax depth as they are likely correlated with the tube length and flower gape. The head, thorax, and abdomen of each pollinator were swabbed for pollen using fuchsin gel, in which the number of pollen grains was counted using a compound microscope and identified as either *G. carneus* or heterospecific pollen, which was determined from pollen controls taken from each population. Pollen loads larger than 1 000 grains on each part of the pollinator were not counted any further. Insects recorded visiting and making contact with the reproductive parts of the flowers or found with *G. carneus* pollen on the head, thorax, or abdomen were considered to be legitimate pollinators and used in the subsequent analyses. Once pollinators were identified, they were classified into functional pollinator groups based on similarities in body size, proboscis length, and foraging behaviours. These functional pollinator groups were used for all subsequent analyses.

A series of networks were used to associate the main functional pollinators for each population and variety. We calculated the visitation rate (visits.flower^-1^.hour^-1^), mean pollen loads and pollinator importance (visitation rate * pollen loads) per functional pollinator for each site (*see Table S5 for sample sizes*). The visitation rates, pollen loads, and pollinator importance were then visualised in a network using the *plotweb* function. For all three networks, H_2_’ was used to assess the overall level of specialisation for the entire network. The H_2_’ values range between 0 (indicating no specialisation) and 1 (indicating complete specialisation) (Bluthgen *et al*., 2006). We used a modularity test on the pollinator importance network to identify different pollinator niches for the *G. carneus* populations. The modularity test was conducted using the *computeModules* and *plotModuleWeb* functions (Beckett, 2016). All of these analyses used the *‘bipartite’* package (Dormann *et al*., 2008).

We further tested whether there were relationships between floral and pollinator morphology using ordinary least squares regressions, specifically between tube length and proboscis length, as well as flower gape and thorax depth. As many sites had more than one functional pollinator, pollinator importance values were used to calculate a ‘weighted’ proboscis length and thorax depth for each site. In the models, the focal floral measurement (either tube length or flower gape) was set as the response, while the pollinator morphology (weighted proboscis length or thorax depth) was set as the explanatory variable. All models were implemented using the *lm* function in the ‘*stats*’ (R Core Team, 2021) package.

### Are differences in the ecological niches of *Gladiolus carneus* varieties resulting in premating reproductive isolation?

#### Calculation of premating barriers

As all the barriers documented affect the co-occurrence of the varieties, we used the first of Sobel and Chen (2014)’s equations to calculate ecogeographic, phenological, and pollinator-mediated isolation.

The equation:

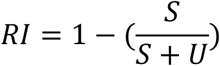

where *S* is the shared and *U* is the unshared portion of occurrence is used to calculate reproductive isolation for prezygotic barriers that effect co-occurrence. The resulting reproductive isolation values range from 0 to 1, where a value of 0 represents complete overlap and absent reproductive isolation, and 1 represents no overlap and complete reproductive isolation.

For ecogeographic isolation, we used the binary niche model predictions of each variety to estimate the shared and unshared predicted niche occupancy for each pair of varieties. Areas where both varieties were present were counted as the ‘shared’ area while areas where only one variety was present was counted as ‘unshared’. When measuring phenological isolation, we were unable to take into account the relative abundances of each flowering variety as suggested by Sobel and Chen (2014) due to differences in sampling efforts on iNaturalist. As varieties such as *macowanianus, albidus, blandus* and *callistus* largely occur in residential areas and on easily accessible hiking routes, they are very well documented. Other varieties such as *high-altitude, langeberg* and *prismatosiphon* largely occur in remote or inaccessible areas, resulting in fewer occurrence points, making it difficult to estimate the number of individuals that are flowering in any area. Therefore, instead of using the relative abundance of each flowering variety, we instead documented binary flowering occurrences on each day of the year to calculate phenological isolation. We used the number of shared and unshared functional pollinators to calculate pollinator-mediated isolation. All legitimate functional pollinators (i.e., insects recorded vising a variety and making contact with the reproductive parts of the flower, or found with *G. carneus* pollen on their head, thorax, or abdomen) were used to calculate pollinator-mediated isolation. As *blandus* lacked any pollinator data, pollinator-mediated isolation was not calculated for any gene flow direction including that variety.

Total reproductive isolation was calculated using equation 4E from Sobel and Chen (2014). The equation calculates total reproductive isolation sequentially from prezygotic barriers that effect co-occurrence, to prezygotic barriers not effecting co-occurrence and finally post-zygotic barriers. As only premating barriers have been included in the calculation, the resulting value represents total premating reproductive isolation. Total premating isolation for gene flow directions involving *blandus* only included ecogeographic and phenological isolation. Total premating isolation for all other gene flow directions were calculated using ecogeographic, phenological, and pollinator-mediated isolation.

## RESULTS

### Are previously and newly described varieties of *Gladiolus carneus* morphologically distinct?

#### Morphological differences between varieties of Gladiolus carneus

The permutation MANOVAs of all morphological traits (*F* = 14.36, *df* = 6, *P* = 0.001, Figure 2), floral traits (*F* = 9.38, *df* = 6, *P* = 0.001, Figure S3A), and vegetative traits (*F* = 17.29, *df* = 6, *P* = 0.001, Figure S3B) showed that the varieties clustered separately. The pairwise MANOVA on all morphological traits found that all pairwise comparisons for the varieties were significant (*P <* 0.05). The pairwise comparisons on the floral traits showed that all pairings were significant (*P <* 0.05), except for *callistus* and *high-altitude* (*P =* 1.00). The pairwise comparisons between varieties based on their vegetative traits showed all pairings were significant (*P <* 0.05), except between *albidus* and *blandus* (*P* = 0.13); and also, *macowanianus* and *callistus* (*P =* 0.19).

**Figure 2.**
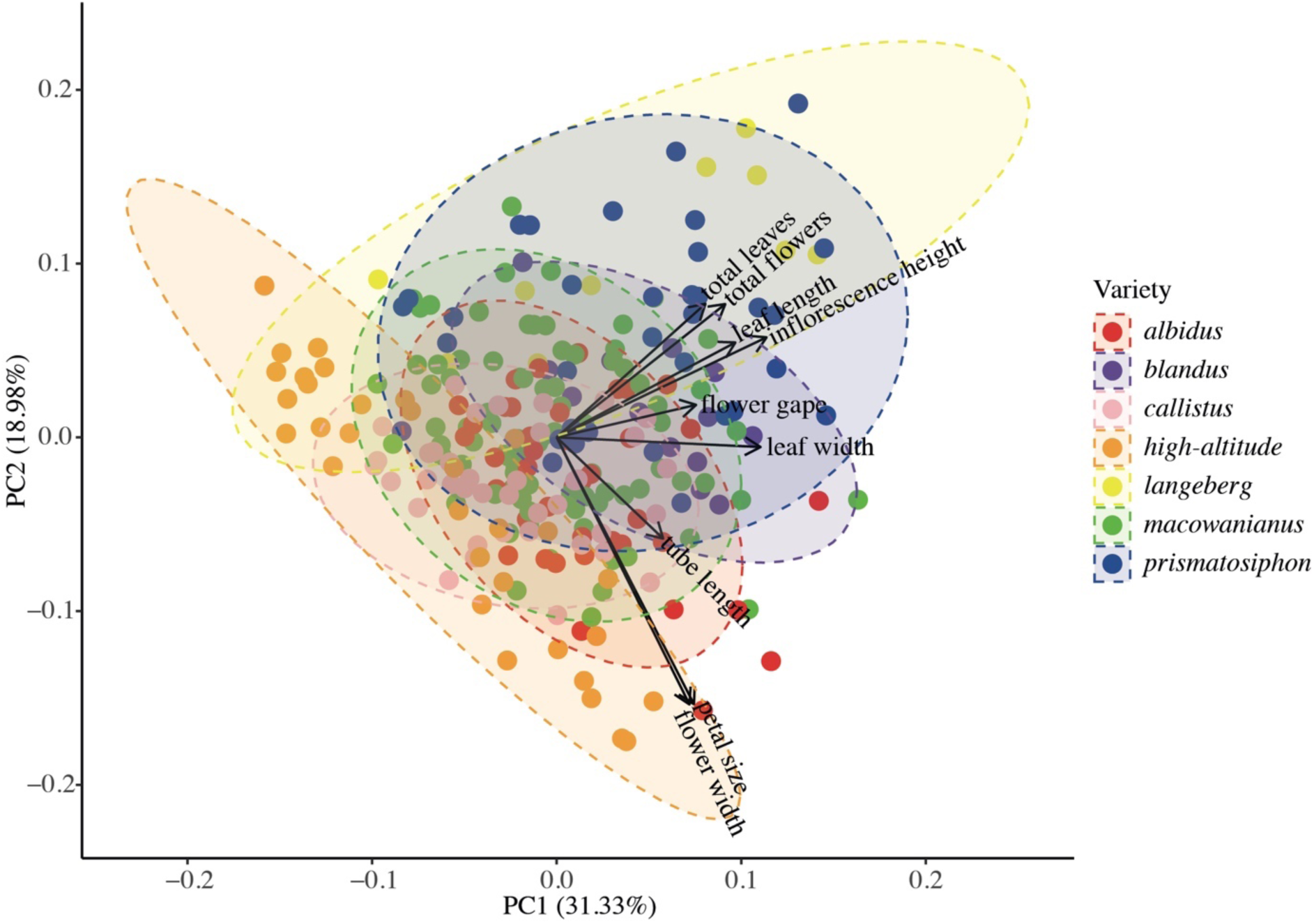
PCA of floral and vegetative traits of *G. carneus* varieties. The PCA includes ellipses showing 95% confidence intervals and a biplot of trait loadings.

We found that both variety and site nested within variety were significant predictors of tube length (*P* < 0.0001, *P* < 0.0001, Table S6, Figure S4A), flower gape (*P* < 0.0001, *P* < 0.0001, Table S6, Figure S4B), petal size (*P* < 0.0001, *P* < 0.0001, Table S6, Figure S4C), inflorescence height (*P* < 0.0001, *P* < 0.0001, Table S6, Figure S4D), the leaf length (*P* < 0.0001, *P* < 0.0001, Table S6, Figure S4E), and leaf width (*P* < 0.0001, *P* < 0.0001, Table S6, Figure S4F).

#### Colour differences between varieties of Gladiolus carneus

There were statistical differences between the varieties for all colour traits modelled in bee and fly colour vision. Specifically, there were significant differences between the varieties’ tepals (Bee: *Approx F* = 39.73, *df* = 6, *P* < 0.0001; Fly: *Approx F* = 38.84, *df* = 6, *P* < 0.0001; Figure 3A, 4A), centre of the median tepals (Bee: *Approx F* = 45.21, *df* = 5, *P* < 0.0001; Fly: *Approx F* = 39.74, *df* = 5, *P* < 0.0001; Figure 3B, 4B), guide of the median tepals (Bee: *Approx F* = 8.57, *df* = 4, *P* < 0.0001; Fly: *Approx F* = 7.68, *df* = 4, *P* < 0.0001; Figure 3C, 4C), centre of the lateral tepals (Bee: *Approx F* = 43.78, *df* = 5, *P* < 0.0001; Fly: *Approx F* = 51.14, *df* = 5, *P* < 0.0001; Figure 3D, 4D), and guide of the lateral tepals (Bee: *Approx F* = 4.54, *df* = 4, *P* < 0.0001; Fly: *Approx F* = 3.58, *df* = 4, *P =* 0.0005; Figure 3E, 4E). Pairwise comparisons showed there were significant differences between the varieties for all of the colour traits within bee and fly colour vision (*see pairwise comparisons in Table S8*). As there were statistical differences between the varieties for all colour traits, it was necessary to test whether those differences are perceptible within bee and fly colour vision.

**Figure 3.**
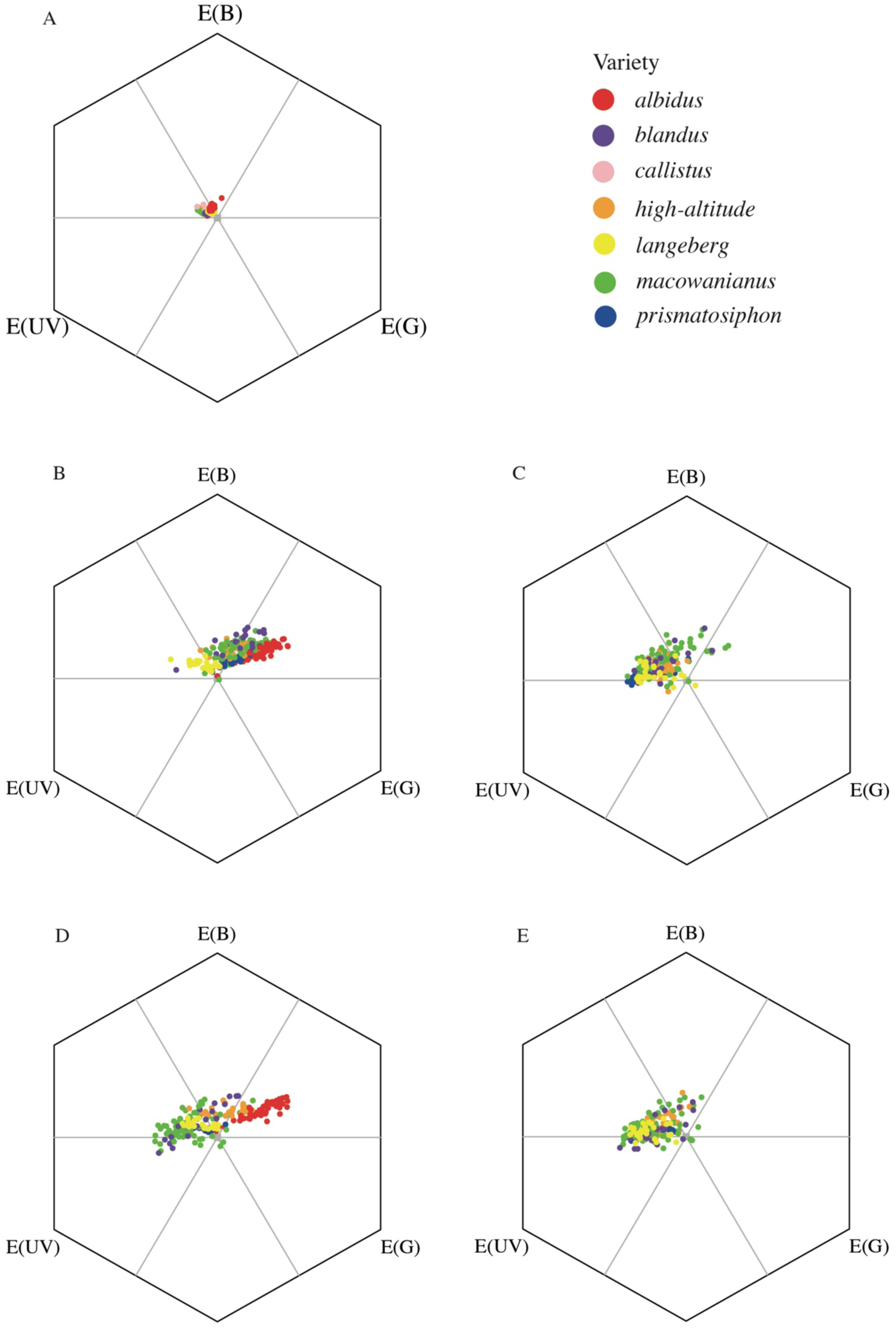
Spectra of *G. carneus* varieties’ (A) tepal, (B) centre of the median tepal, (C) guide of median tepal, (D) centre of lateral tepal, (E) and guide of lateral tepal plotted in bee colour vision with *Apis mellifera* spectral sensitivities.

**Figure 4.**
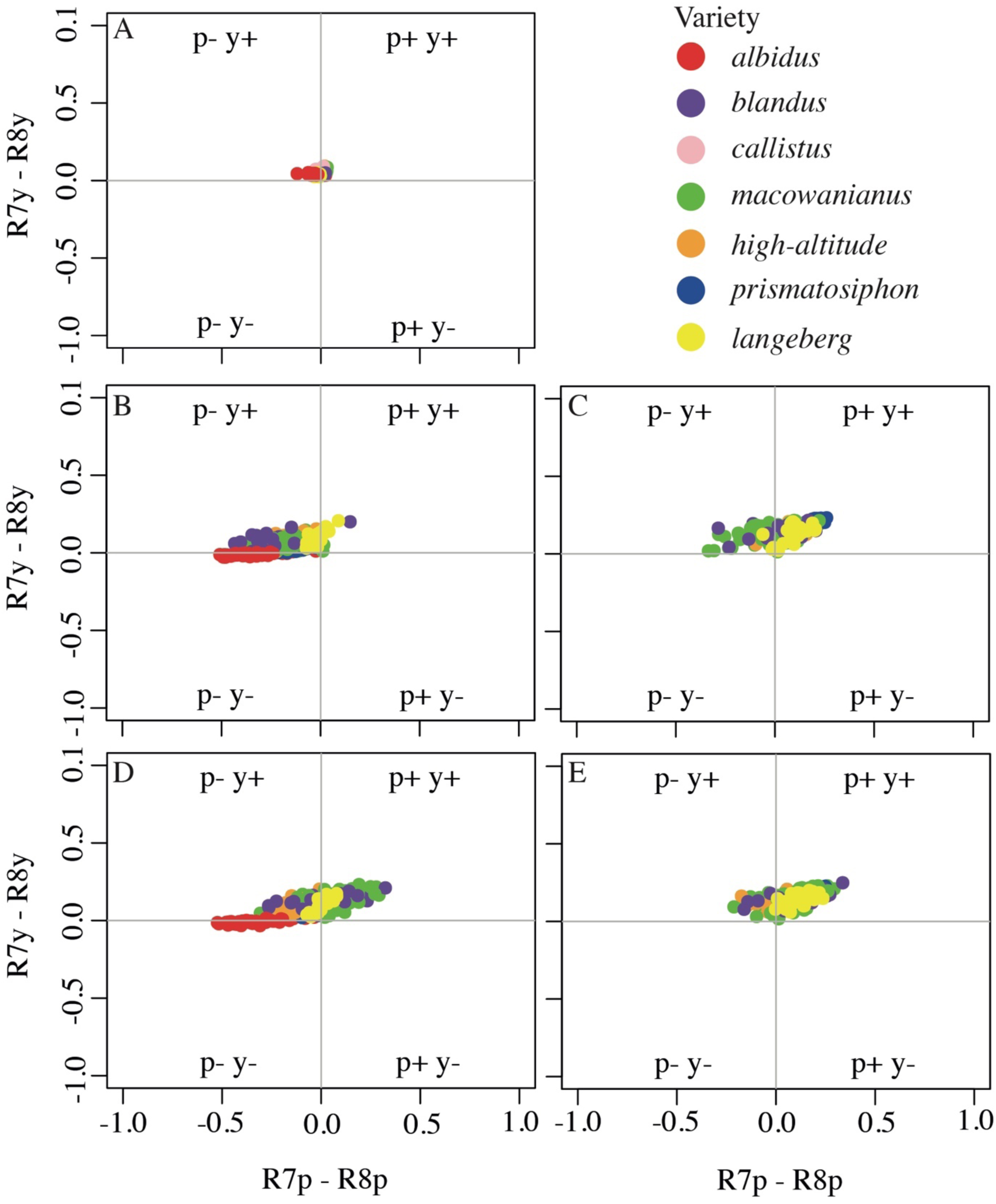
Spectra of *G. carneus* varieties’ (A) tepal, (B) centre of the median tepal, (C) guide of median tepal, (D) centre of lateral tepal, (E) and guide of lateral tepal plotted in fly colour vision with *Eristalis tenax* spectral sensitivities.

All comparisons between the varieties’ tepals were below the perceptibility threshold within the bee and fly colour vision models suggesting that bees or flies would not be able to distinguish between the varieties tepals (Figure S5A, S6A). As *callistus* did not have any nectar guides, the variety has been left out of all further comparisons. Similarly, only comparisons with the centre of *albidus* guides were included as they lack any outline of the nectar guides. For the centre of the lower median tepal, modelled in bee and fly vision, there are perceptible differences between *langeberg* and all other varieties, and between *albidus* and all varieties except *macowanianus* (Figure S5B, S6B). Additionally, the comparisons between *albidus* and *macowanianus*; and *blandus* and *prismatosiphon* were only perceptible within fly vision (Figure S6B). For the centres of the lower lateral tepals, there are perceptible differences between *albidus* and all varieties and *high-altitude* and all varieties in bee and fly colour vision (Figure S5D, S6D). The contrast between *macowanianus* and *prismatosiphon* is also perceptible within bee and fly vision (Figure S5D, S6D). Within fly vision, the comparison between *blandus* and *prismatosiphon* was also perceptible (Figure S6D). For the guide of the lower median tepal, the contrast between *macowanianus* and *prismatosiphon* were perceptible within bee and fly colour vision (Figure S5C, S6C). The contrasts between the guide of the lower median tepal was perceptible between *langeberg* and *macowanianus* and between *high-altitude* and *prismatosiphon* within fly vision (Figure S6C). There were no perceptible contrasts between the guide on the lower lateral tepal within bee vision (Figure S5E), but there were perceptible differences between *high-altitude* and *langeberg* within fly vision (Figure S6E).

### Do *Gladiolus carneus* varieties occupy distinct biotic and abiotic niches?

#### Do varieties of Gladiolus carneus occupy distinct abiotic niches?

The MaxEnt models for all varieties performed well (AUC > 0.95), providing reliable predicted abiotic niches. All species distribution models closely matched the documented distributions of each variety. Specifically, the *albidus* and *callistus* varieties had similar distributions around the lowlands of the Boland (Figure 5A, 5C). *Blandus* had the smallest distribution and was largely restricted to the Southern Cape coast between Rooiels and Hermanus (Figure 5B). The *high-altitude* variety was restricted to the Drakenstein, Hottentots-holland and Riviersonderend mountain ranges (Figure 5D). The *langeberg* variety was predicted to occur throughout the Langeberg Mountain range (Figure 5E). *Macowanianus* had a predicted range covering the peninsula and Southern Cape coast (Figure 5F), whilst *prismatosiphon* was largely restricted to the Southern Overberg (Figure 5G). The jackknife tests indicated that annual precipitation (mm) was the most important variable in the *albidus, callistus,* and *high-altitude* models, the annual temperature range (°C) was most important for the *blandus* and *macowaninaus* models, precipitation seasonality was the most important for the *langeberg* model and extractable phosphorus (mg/kg) was the most important for the *prismatosiphon* model. The permutation MANOVA showed that there were significant differences in the realised abiotic niches of the varieties (*F* = 100.29, *df* = 6, *P* = 0.001, Figure 6). Furthermore, the pairwise permutation MANOVA indicated that all varieties occupied realised abiotic niches that were significantly different from each other (*P* < 0.05).

**Figure 5.**
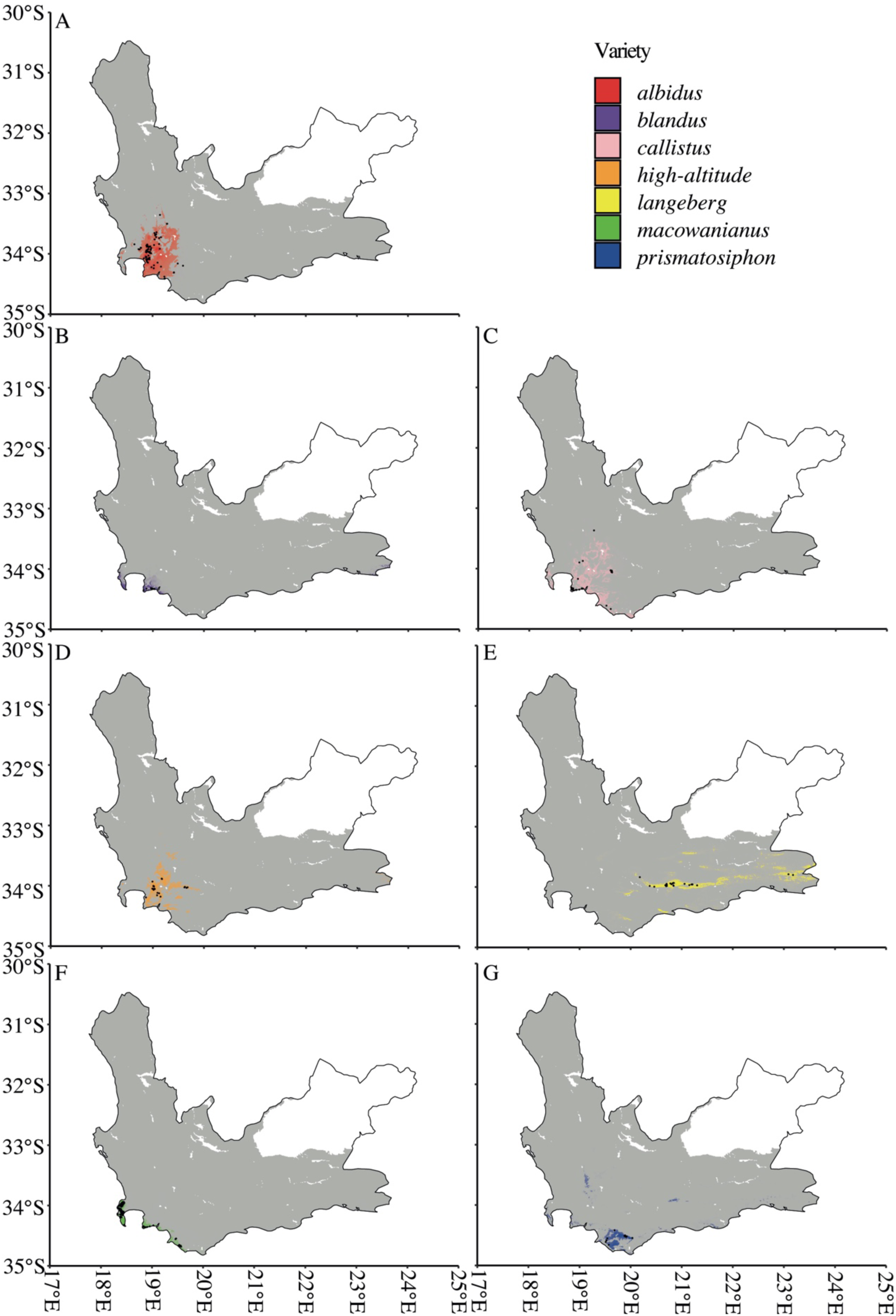
Niche model outputs for (A) *albidus*, (B) *blandus*, (C) *callistus*, (D) *high-altitude*, (E) *langeberg*, (F) *macowanianus*, and (G) *prismatosiphon* based on uncorrelated bioclimatic, elevation and soil layers. Each map shows the Western Cape of South Africa. The grey area is the extent of the modelled area, coloured areas show the model predictions, and black points show localities.

**Figure 6.**
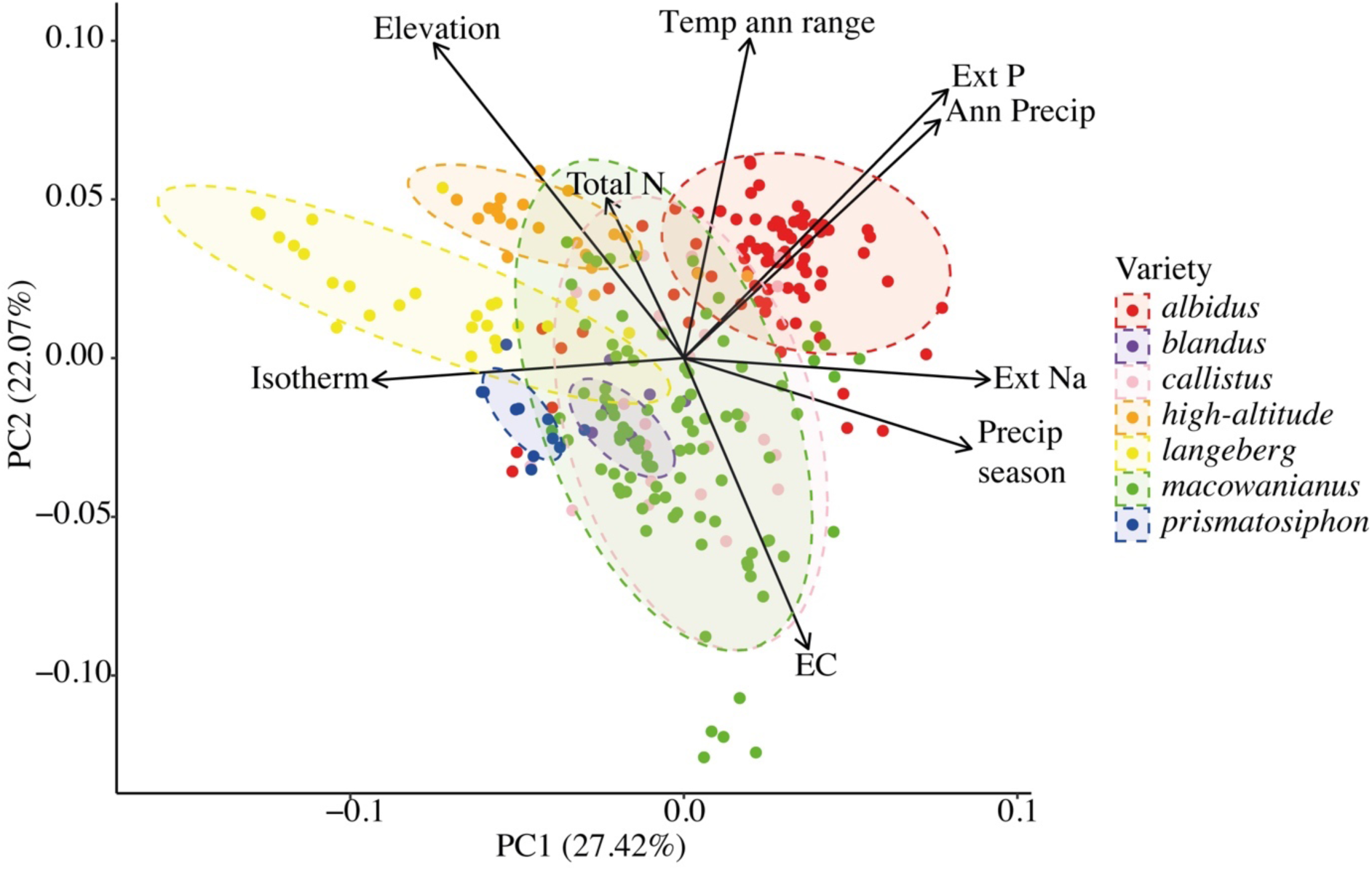
PCA of the abiotic niche of the *G. carneus* varieties. Abiotic layers include uncorrelated bioclimatic, elevation and soil variables. The variables included are: elevation, isothermality, annual temperature range, annual precipitation, precipitation seasonality, electrical conductivity, total nitrogen, extractable phosphorus, and extractable sodium.

#### Do varieties of Gladiolus carneus flower at different times?

There were significant differences in the flowering times between *G. carneus* varieties (*W* = 389.23, *df* = 12, *P* < 0.0001; Figure 7), suggesting that different varieties occupy distinct phenological niches. The pairwise comparisons between varieties flowering times showed all pairings were significant (*P* < 0.01) except between *blandus* and *langeberg* (*P* = 0.10), and *callistus* and *prismatosiphon* (*P* = 1.00).

**Figure 7.**
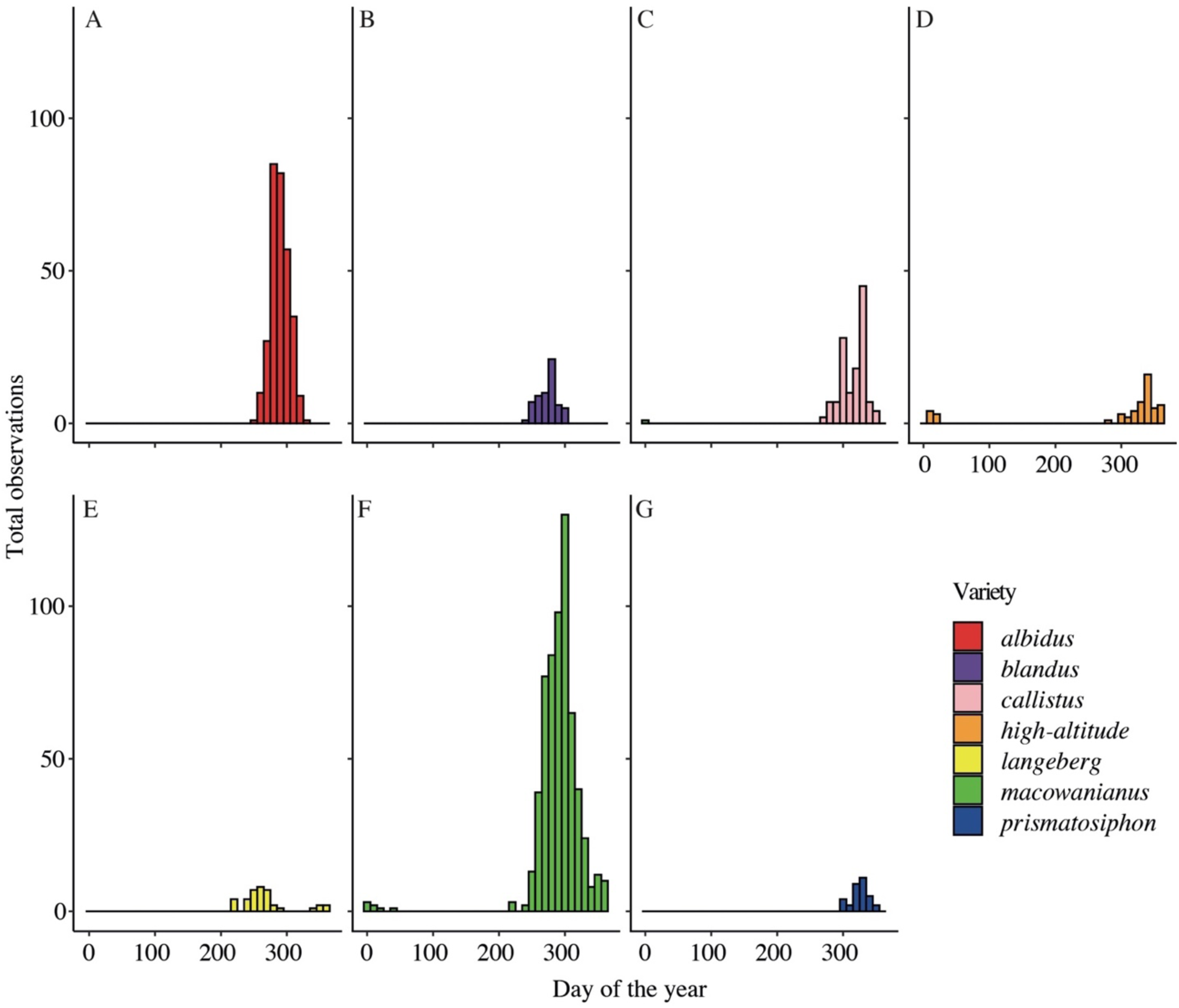
Flowering phenology of *G. carneus* varieties shown by the total iNaturalist observations on each day of the year. *G. carneus* varieties are (A) *albidus*, (B) *blandus*, (C) *callistus*, (D) *high*-*altitude*, (E) *langeberg*, (F) *macowanianus*, and (G) *prismatosiphon*.

#### Do varieties of Gladiolus carneus occupy distinct pollinator niches?

Across all sampled *G. carneus* populations, a diverse range of pollinators were found which were subsequently classified into the following functional groups: solitary bees, carpenter bees, long-tongued flies (LTFs), medium-tongued flies (MTFs), honey bees and Lycaenid butterflies. Solitary bees included *Amegilla spilostoma, Amegilla obscuritarsis,* and *Anthophora diversipes*; carpenter bees included all species belonging to the genus *Xylocopa*; LTFs included *Philoliche rostrata* and *Moegistorhynchus manningi* which had proboscis between 28 and 39mm; MTFs included *Prosoeca nitidula*, *Prosoeca westermanii*, and *Philoliche lateralis* which had proboscis between 4 and 16mm; honey bees included all *Apis mellifera* individuals.

The networks of visitation rates (*H*2’ = 0.76, Figure 8A), pollen loads (*H*2’ = 0.71, Figure 8B) and pollinator importance (*H*2’ = 0.99, Figure 8C) all showed high levels of specialisation. In particular, the pollinator importance network showed near complete specialisation with each population having a single highly effective functional pollinator. Across all *G. carneus* populations, there were only three important functional pollinator groups: solitary bees, MTFs and LTFs. The modularity analysis on the pollinator importance network showed that *G. carneus* populations were associated with the same pollinator niches: solitary bees, MTFs and LTFs (Figure S7). Although the network and modularity analysis showed high levels of specialisation to a specific functional pollinator at the population level, at the variety level, the most important functional pollinator was not consistent. The exception to this trend was *albidus*, which was consistently pollinated by solitary bees across all sampled populations.

**Figure 8.**
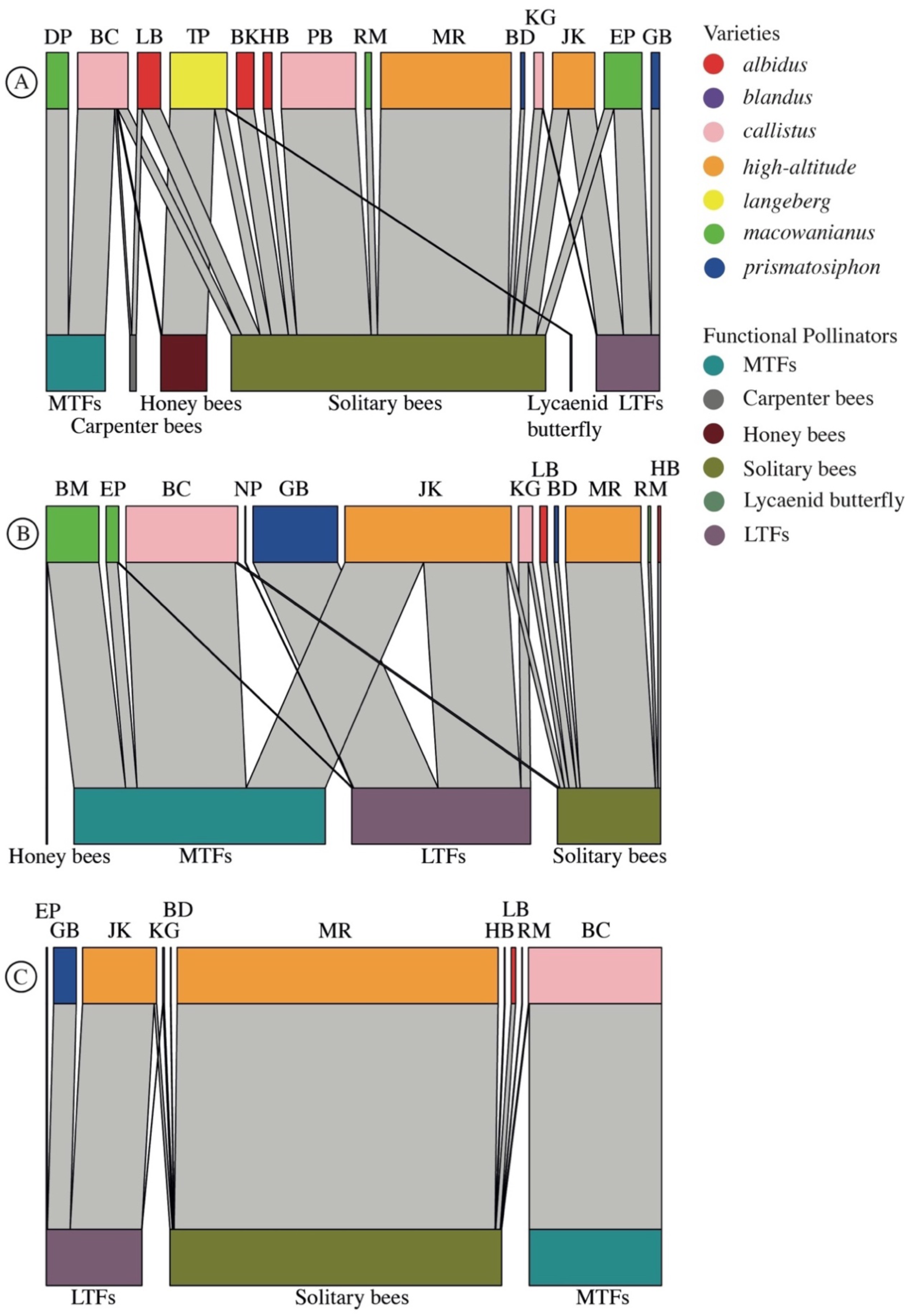
Networks of the (A) visitation rates, (B) pollen loads, and (C) pollinator importance of the functional pollinator groups: solitary bees, honey bees, carpenter bees, medium-tongued flies (MTFs), long-tongued flies (LTFs), and Lycaenid butterflies to different populations of *G. carneus* varieties.

*Gladiolus carneus* populations show high levels of specialisation towards a single functional pollinator, and we further found a significant positive relationship between the tube length and weighted proboscis length (*R*^2^ = 0.46, *F* = 8.622, *df* = 8, *P* = 0.02; Figure S8A); however, there was no significant relationship between flower gape and weighted thorax depth (*R*^2^ = 0.01, *F* = 1.10, *df* = 8, *P* = 0.33; Figure S8B).

### Are differences in the ecological niches of *Gladiolus carneus* varieties resulting in premating reproductive isolation?

Ecogeographic isolation was the strongest gene flow barrier across the *G. carneus* species complex (RI_ecogeo_ = 0.84 ± 0.03SE, Table 1). There was weak ecogeographic isolation from *albidus* to *callistus* and the *high-altitude* variety (Ecogeo: RI_al → cl_ = 0.36, RI_al → ha_ = 0.45), and from *callistus* to *macowanianus* (Ecogeo: RI_cl → mc_ = 0.54). There was also weak ecogeographic isolation from *albidus, callistus,* and *macowanianus* to *blandus* (Ecogeo: RI_al → bl_ = 0.45, RI_cl → bl_ = 0.40, RI_mc → bl_ = 0.40). The small range of *blandus* is largely encased within the larger ranges of *albidus*, *callistus* and *macowanianus,* resulting in asymmetric ecogeographic isolation. All other gene flow directions between the varieties were above 0.6 indicating strong ecogeographic isolation, with many gene flow directions being near complete (Table 1).

**Table 1.** Ecogeographic, phenological, pollinator-mediated isolation and total premating isolation for all gene flow directions between the *G. carneus* varieties. There are no estimates of pollinator-mediated isolation for gene flow directions including *blandus* due to a lack of pollinator data. Total premating isolation has been estimated using ecogeographic and phenological isolation for gene flow directions including *blandus*. Premating isolation has been calculated using ecogeographic, phenological and pollinator-mediated isolation for all other gene flow directions.

| Gene flow direction | Ecogeographic isolation | Phenological isolation | Pollinator-mediated isolation | Total premating isolation |
| --- | --- | --- | --- | --- |
| *albidus* -> *blandus* | 0.45 | 0.27 | - | 0.60 |
| *albidus* -> *callistus* | 0.36 | 0.38 | 0.60 | 0.84 |
| *albidus* -> *high-altitude* | 0.45 | 0.88 | 0.67 | 0.98 |
| *albidus* -> *langeberg* | 1.00 | 0.72 | 0.67 | 1.00 |
| *albidus* -> *macowanianus* | 0.73 | 0.46 | 0.75 | 0.96 |
| *albidus* -> *prismatosiphon* | 0.93 | 0.58 | 0.50 | 0.99 |
| *blandus* -> *albidus* | 0.96 | 0.59 | - | 0.98 |
| *blandus* -> *callistus* | 0.93 | 0.83 | - | 0.99 |
| *blandus* -> *high-altitude* | 0.97 | 0.94 | - | 1.00 |
| *blandus* -> *langeberg* | 1.00 | 0.59 | - | 1.00 |
| *blandus* -> *macowanianus* | 0.75 | 0.73 | - | 0.93 |
| *blandus* -> *prismatosiphon* | 0.96 | 1.00 | - | 1.00 |
| *callistus* -> *albidus* | 0.61 | 0.51 | 0.00 | 0.81 |
| *callistus* -> *blandus* | 0.40 | 0.76 | - | 0.85 |
| *callistus* -> *high-altitude* | 0.76 | 0.69 | 0.00 | 0.93 |
| *callistus* -> *langeberg* | 1.00 | 0.91 | 0.33 | 1.00 |
| *callistus* -> *macowanianus* | 0.54 | 0.59 | 0.00 | 0.81 |
| *callistus* -> *prismatosiphon* | 0.95 | 0.21 | 0.00 | 0.96 |
| *high-altitude* -> *albidus* | 0.70 | 0.93 | 0.50 | 0.99 |
| *high-altitude* -> *blandus* | 0.80 | 0.94 | - | 0.99 |
| *high-altitude* -> *callistus* | 0.78 | 0.79 | 0.40 | 0.97 |
| *high-altitude* -> *langeberg* | 1.00 | 0.94 | 0.67 | 1.00 |
| *high-altitude* -> *macowanianus* | 0.91 | 0.82 | 0.25 | 0.99 |
| *high-altitude* -> *prismatosiphon* | 0.91 | 0.79 | 0.00 | 0.98 |
| *langeberg* -> *albidus* | 1.00 | 0.85 | 0.50 | 1.00 |
| *langeberg* -> *blandus* | 1.00 | 0.61 | - | 1.00 |
| *langeberg* -> *callistus* | 1.00 | 0.94 | 0.60 | 1.00 |
| *langeberg* -> *high-altitude* | 1.00 | 0.94 | 0.67 | 1.00 |
| *langeberg* -> *macowanianus* | 1.00 | 0.80 | 0.50 | 1.00 |
| *langeberg* -> *prismatosiphon* | 0.92 | 0.89 | 0.50 | 1.00 |
| *macowanianus* -> *albidus* | 0.95 | 0.02 | 0.50 | 0.98 |
| *macowanianus* -> *blandus* | 0.40 | 0.12 | - | 0.47 |
| *macowanianus* -> *callistus* | 0.87 | 0.06 | 0.20 | 0.90 |
| *macowanianus* -> *high-altitude* | 0.97 | 0.41 | 0.00 | 0.98 |
| *macowanianus* -> *langeberg* | 1.00 | 0.34 | 0.33 | 1.00 |
| *macowanianus* -> *prismatosiphon* | 0.94 | 0.11 | 0.00 | 0.94 |
| *prismatosiphon* -> *albidus* | 0.98 | 0.86 | 0.50 | 1.00 |
| *prismatosiphon* -> *blandus* | 0.87 | 1.00 | - | 1.00 |
| *prismatosiphon* -> *callistus* | 0.98 | 0.68 | 0.60 | 1.00 |
| *prismatosiphon* -> *high-altitude* | 0.96 | 0.88 | 0.33 | 1.00 |
| *prismatosiphon* -> *langeberg* | 0.96 | 0.94 | 0.67 | 1.00 |
| *prismatosiphon* -> *macowanianus* | 0.91 | 0.84 | 0.50 | 0.99 |
| **Mean reproductive isolation** | **0.85 ± 0.03SE** | **0.67 ± 0.04SE** | **0.39 ± 0.04SE** | **0.95 ± 0.02SE** |

Phenological isolation was a relatively strong gene flow barrier across the species complex (RI_phenology_ = 0.67 ± 0.04SE, Table 1). There was weak phenological isolation from *albidus* to *callistus* (Phenology: RI_al → cl_ = 0.27) and from *callistus* to *prismatosiphon* (Phenology: RI_cl → pr_ = 0.21) due to overlapping flowering times. There was also weak phenological isolation from *macowanianus* to *albidus, blandus, callistus* and *prismatosiphon* (Phenology: RI_mc → al_ = 0.02, RI_mc → bl_ = 0.12, RI_mc → cl_ = 0.06, RI_mc → pr_ = 0.11). *Macowanianus* has the longest flowering time (stretching from September to early January) resulting in a large overlap in the flowering times of other varieties, allowing for gene flow in one direction, but not the other.

Pollinator-mediated isolation was a relatively weak gene flow barrier between the varieties (RI_pollinator_ = 0.39 ± 0.05SE, Table 1). Pollinator-mediated isolation was weakest from *callistus* to *albidus, high-altitude, macowanianus* and *prismatosiphon* (Pollinator: RI_cl → al_ = 0.00, RI_cl → ha_ = 0.00, RI_cl → mc_ = 0.00, RI_cl → pr_ = 0.00), and from *high-altitude* to *prismatosiphon* and *macowanianus* (Pollinator: RI_ha → pr_ = 0.00, RI_ha → mc_ = 0.25), and from *macowanianus* to *prismatosiphon*, *high-altitude* and *callistus* (Pollinator: RI_mc → pr_ = 0.20, RI_mc → ha_ = 0.00, RI_mc → cl_ = 0.00). *Callistus, macowanianus* and *high-altitude* varieties have broad pollinator niches resulting in a high potential gene flow to other varieties.

The combined effects of ecogeographic, phenological and pollinator-mediated isolation has resulted in near complete premating isolation (RI = 0.95 ± 0.02SE, Table 1) across the species complex. The only weak gene flow direction for total premating reproductive isolation was from *macowanianus* to *blandus* (RI_mc → bl_ = 0.47).

## DISCUSSION

We provide evidence that morphologically distinct varieties of *G. carneus* occupy distinct realized and fundamental abiotic niches resulting in strong ecogeographic isolation across the species complex. The varieties additionally have different flowering times that cause moderate phenological isolation. Although individual populations of *G. carneus* were pollinated by a single, highly effective functional pollinator, at the level of the variety, plants had multiple highly effective functional pollinators resulting in relatively weak pollinator-mediated isolation. The strength of these premating barriers, and in particular ecogeographic isolation, caused near complete reproductive isolation between the varieties. These results suggest that niche differentiation, and particularly abiotic factors, may be playing a role in driving incipient speciation within the species complex.

### Are Gladiolus carneus varieties morphologically distinct?

All varieties showed evidence of being morphologically distinct in both floral and vegetative traits lending support to the varieties initially described by Delpierre and du Plessis (1974) and for the two newly described varieties, *langeberg* and the *high-altitude* varieties. The varieties also differ significantly in their colour traits. *Albidus* only has a large central UV absorbent nectar guide (Figure S1E) and *callistus* has no nectar guides and instead has a dark purple gullet (Figure S1F). All other varieties have distinct centres and guides on their lower tepals (Figure S1D). The bee and fly colour vision models indicated that most of the perceptible differences between the varieties were in the centres of the median and lateral tepals. In particular, *langeberg* and *albidus* had distinct centres on their median tepals and *albidus*, *high-altitude* and *prismatosiphon* had distinct centres on their lateral tepals. These divergent colour properties are likely associated with adaptations to distinct functional pollinators (Demarche *et al*., 2015). In particular, the UV absorbent centres of *albidus* have likely evolved to attract their pollinators, solitary bees (Chittka, 1992, Demarche *et al*., 2015, de Camargo *et al*., 2019, Finnell and Koski, 2021, Newman *et al*., 2022). The purple and red nectar guides found on all other varieties, except for *callistus,* is consistent with nectar guides documented in long proboscid Nemestrinid and Philoliche fly systems, and may represent a unique adaptation to the long proboscid fly pollination syndrome (Goldblatt *et al*., 2001, Newman *et al*., 2014). However, since the varieties are not associated with a single highly effective functional pollinator, direct associations between nectar guide properties and functional pollinators should be assessed at the population level with reference to the most important pollinator (Moir *et al*., 2025).

### Abiotic niche differentiation causing ecogeographic isolation

*Gladiolus carneus* varieties occupied distinct realised and fundamental niches (Figure 5, 6) leading to strong ecogeographic isolation across the species complex. There were a number of cases of asymmetric ecogeographic isolation, firstly caused by the large *albidus* range overlapping with the *callistus* and *high-altitude* varieties, and secondly from the small *blandus* range being largely encased within the ranges of *albidus, callistus* and *macowaniaus.* Across plant taxa, ecogeographic isolation is relatively symmetrical, however individual cases can vary substantially, which largely corroborates the results found in the *G. carneus* complex (Christie *et al*., 2022). These results, along with the differences in vegetative morphology documented between the varieties (Figure S2B), suggests that abiotic niche differentiation plays a potential role in the diversification of the species complex. The importance of abiotic niche differentiation driving diversification is largely congruent with previous experimental evidence in the CFR (e.g., Latimer *et al*., 2009, Carlson *et al*., 2011, Mitchell *et al*., 2015, Carlson *et al*., 2016). For example, Carlson *et al*. (2011) found that white *Protea* populations differ in functional traits across environmental gradients, some of which are likely maintained by divergent selection imposed by ecological differences. Macroevolutionary evidence supporting abiotic niche differentiation in the CFR is somewhat mixed. van der Niet and Johnson (2009) tested for ecological shifts in 188 Cape sister species and found evidence that habitat, pollinator and fire-frequency shifts were more frequent than soil type shifts, while Schnitzler *et al*. (2011) analysed 470 species from the phylogenies of four Cape clades and found that soil type had the highest variability between sister species in three of the clades. Outside of the CFR, abiotic niche differentiation is a major driver of diversification (Grossenbacher *et al*., 2014, Fernandez-Mazuecos and Glover, 2024) and the resulting gene flow barrier, ecogeographic isolation, has been found to be one of the strongest gene flow barriers between recently diverged taxa (Christie *et al*., 2022).

### Flowering time differences causing phenological isolation

There were differences in the flowering times of the *G. carneus* varieties leading to varying strengths of phenological isolation (Figure 7). In a few cases, phenological isolation was asymmetric which was largely due to *macowanianus* having a long flowering time that overlapped with *albidus, blandus, callistus* and *prismatosiphon*. These shifts in phenology may also be driving the diversification within the species complex. Phenological shifts have been well documented within the CFR, with van der Niet and Johnson (2009) showing that over 30% of 188 species pairs from eight Cape clades had a flowering time shift. Similarly, Linder (2020) found that 47% of 112 sister species pairs in the Restionaceae had non-overlapping flowering times. These findings indicate that phenological shifts seem to be a relatively frequent isolating barrier between taxa in the CFR (Ellis *et al*., 2014). However, this seems to contradict the results from a global comparison showing phenological isolation seems to play a lessor role in speciation events between recently diverged taxa (RI_phenology_ = 0.38) (Christie *et al*., 2022).

### Differing functional pollinators causing pollinator-mediated isolation

Although each population of *G. carneus* had a single highly effective functional pollinator, each variety, apart from *albidus,* was not associated a single functional pollinator. This suggests that the varieties do not occupy distinct pollination niches and that pollinators are unlikely to be driving the diversification at the variety level. These results are similar to Schnitzler *et al*. (2011) which found that in four highly diverse Cape Clades, pollinator shifts were relatively infrequent compared to shifts in soil type and fire survival strategy. However, these results contrast with both phylogenetic and experimental evidence showing that pollinator-driven speciation is common in both the CFR (van der Niet and Johnson, 2009, Johnson, 2010, Anderson *et al*., 2014, Forest *et al*., 2014, Newman *et al*., 2015, Newman and Johnson, 2021) and in the genus *Gladiolus* (Anderson *et al*., 2010, Valente *et al*., 2012). Furthermore, Christie *et al*. (2022) found pollinator-mediated isolation, along with ecogeographic isolation, was one of the strongest reproductive isolation barriers documented across 89 taxa pairs of seed plants. However, as populations of *G. carneus* only had one highly effective functional pollinator and there were associations between floral and pollinator morphology across sites, pollinators may be driving population level shifts and causing reproductive isolation between populations rather than between varieties. Alternatively, the estimates of pollinator-mediated isolation may be inaccurate. Taking into account only the shared and unshared functional pollinators does not account for differential efficiency of functional pollinators to alternative varieties (Ostevik *et al*., 2016, Moir *et al*., 2025). The strength of pollinator-mediated isolation could be more accurately estimated between varieties with sympatric populations (e.g., *macowanianus* and *callistus*) where pollinator transitions (Sobel and Streisfeld, 2015) and pollen flow (Kay, 2006) could be used to estimate behavioural and mechanical isolation.

### Should Gladiolus carneus varieties be treated as separate taxa?

The biological species concept posits that taxa should be treated as separate species when they are reproductively isolated from one another (Coyne and Orr, 2004). All varieties of *G. carneus* showed evidence of distinct vegetative and floral morphology (Figure 1, 2) and are strongly reproductively isolated from one another (RI_total_ = 0.95 ± 0.02SE). The only exception, *blandus*, differs only in vegetative morphology from *macowanianus* and is encased within the geographic range and flowering time of *macowanianus,* causing weak reproductive isolation. As premating reproductive isolation is near compete between the varieties, they can likely be treated as separate taxa regardless of strength of postpollination isolation. In the CFR, there are examples of interfertile orchids being treated as separate taxa (Newman and Johnson, 2024). However, whether these taxa should be treated as separate species depends on whether species are defined as being reproductively isolated or when reproductive isolation is irreversible (Lowry, 2012). If species are defined by whether they are reproductively isolated from one another, premating barriers alone can be used to delineate species. However, divergent selection on traits that mediate reproductive isolation can allow for introgression and result in the collapse of species boundaries in the future (Lowry, 2012). If species are instead defined by whether reproductive isolation is irreversible, separate species would only be delineated when intrinsic postzygotic isolation is complete. The *G. carneus* varieties, apart from *blandus*, meet the criteria of being reproductively isolated from one another, but it remains to be seen if it is reversable. Further research should quantify the extent of postpollination isolation between the varieties and determine its contribution to total isolation.

## CONCLUSION

Overall, there is evidence of seven morphologically distinct varieties within the *G. carneus* species complex that occupy contrasting abiotic and biotic niches resulting in near complete premating reproductive isolation. These results suggest niche differentiation likely plays a key role in driving the diversification both within this species complex and within the CFR. Future research should test for local adaptation using reciprocal translocations and disentangle the specific factors causing both morphological and niche differentiation between closely related taxa. Additionally, postpollination isolation should be quantified between taxa in the CFR to determine both its relative contribution to maintaining reproductive isolation, and whether reproductive isolation is reversable.

## SUPPLEMENTARY DATA

Supplementary data are available at Annals of Botany online and consist of the following: Table S1. Coordinates, and elevation (m) of all 29 *G. carneus* sites sampled for this manuscript. Table S2. Sample sizes of the morphological trait measurements at each *G. carneus* study population. Table S3. Sample sizes of individuals for which spectral reflectance were collected for each *G. carneus* study population and variety. Table S4. Bioclimatic and topography layers mined from Worldclim and the soil layers from Cramer *et al*. (2019) that were used in the abiotic niche analysis. Table S5. Mean and sample size of visitation rates and pollen loads for each functional pollinator collected across populations and varieties of *G. carneus.* Table S6. Generalised linear model outputs testing for differences between the *G. carneus* varieties morphological traits. Table S7. Pairwise comparisons between the morphological traits of *G. carneus* varieties. Table S8. Pairwise comparisons between varieties of *G. carneus* plotted in bee and fly colour vision models. Figure S1. Morphological measurements of *G. carneus* varieties. Figure S2. Spectral reflectance curves of (A) tepal, (B) gullet, (C) centre of the median tepal, (D) guide of median tepal, (E) centre of lateral tepal, (F) guide of lateral tepal of all *G. carneus* varieties. Figure S3. PCAs of (A) floral and (B) vegetative traits of the *G. carneus* varieties. Figure S4. Comparisons of the functional traits (A) tube length, (B) flower gape, (C) petal size, (D) inflorescence height, (E) leaf length and (F) leaf width between *G. carneus* varieties. Figure S5. Euclidean distances between each variety for the (A) tepal, (B) centre of the median tepal, (C) guide of median tepal, (D) centre of lateral tepal, (E) and guide of lateral tepal modelled in bee colour vision with *Apis mellifera* spectral sensitivities. Figure S6. Euclidean distances between each variety for (A) tepal, (B) centre of the median tepal, (C) guide of median tepal, (D) centre of lateral tepal, (E) and guide of lateral tepal modelled in fly colour vision with *Eristalis tenax* spectral sensitivities. Figure S7. Modularity analysis of pollinator importance showing *G. carneus* sites associated with each functional pollinator group. Figure S8. Correlations between the (A) weighted proboscis length (mm) of pollinators and *G. carneus* tube length (mean ± SE mm), and (B) weighted thorax depth (mm) of pollinators and *G. carneus* flower gape (mean ± SE mm) across populations of *G. carneus*.

## FUNDING

This research was financially supported by the Botanical Education Trust (KLK) and NRF-Thuthuka Grant (ELN) (grant number: TTK210211585733).

## CONFLICTS OF INTEREST

None declared.

## AUTHOR CONTRIBUTIONS

EN and KLK conceptualised the study. KLK and ELN collected the data. KLK analysed all data, and wrote the paper with input from ELN and SE. All authors provided input on editing the manuscript.

## ACKNOWLEDGEMENTS

We thank Kaelin du Plessis, Mallory Hansford, Cindy Fourie, and Marelise Faul for assistance with data collection. Adrian Dyer for providing us with *Eristalis tenax* spectral sensitivities and Hugo Gruson for assisting K.K. with visual modelling. Connal Eardley, for providing bee identifications. Bridgette McMillan, for assisting with the environmental niche modelling. Paula Strauss, for arranging access to Farm 215 and Lesley Richardson and Stephen Smuts for providing access to the Napier mountain conservancy. Robyn Khoury, Tafadzwa Thabethe, Josie Makkink, and the Hansford family for providing accommodation. We also thank Cape Nature (CN35-87-18949; CN44-87-18954) and SANPARKS (CRC/2022-2023/018--2021/V1) for providing permits to conduct this research.

## Notes

### Competing Interest Statement

The authors have declared no competing interest.

## LITERATURE CITED

Agostinelli C, Lund U. 2024. R package ‘circular’: circular statistics. version 0.5-1. https://CRAN.R-project.org/package=circular.

Anderson B, Alexandersson R, Johnson SD. 2010. Evolution and coexistence of pollination ecotypes in an African Gladiolus (Iridaceae). Evolution, 64: 960–972.

Anderson B, Ros P, Wiese TJ, Ellis AG. 2014. Intraspecific divergence and convergence of floral tube length in specialized pollination interactions. Proceedings of the Royal Society B: Biological Sciences, 281: 20141420.

Beckett SJ. 2016. Improved community detection in weighted bipartite networks. Royal Society Open Science, 3: 140536.

Bluthgen N, Menzel F, Bluthgen N. 2006. Measuring specialization in species interaction networks. BMC Ecology, 6: 9.

Boucher FC, Verboom GA, Gallien L, Ellis AG. 2023. Multiple reproductive barriers maintain species boundaries in stone plants of the genus Argyroderma. Botanical Journal of the Linnean Society, 204: 187–197.

Briscoe AD, Chittka L. 2001. The evolution of colour vision in insects. Annual Review of Entomology, 46: 471–510.

Carlson JE, Adams CA, Holsinger KE. 2016. Intraspecific variation in stomatal traits, leaf traits and physiology reflects adaptation along aridity gradients in a South African shrub. Annals of Botany, 117: 195–207.

Carlson JE, Holsinger KE, Prunier R. 2011. Plant responses to climate in the Cape Floristic Region of South Africa: evidence for adaptive differentiation in the Proteaceae. Evolution, 65: 108–124.

Chittka L. 1992. The colour hexagon: a chromaticity diagram based on photoreceptor excitations as a generalized representation of colour opponency. Journal of Comparative Physiology A, 170: 533–543.

Christie K, Fraser LS, Lowry DB. 2022. The strength of reproductive isolating barriers in seed plants: insights from studies quantifying premating and postmating reproductive barriers over the past 15 years. Evolution, 76: 2228–2243.

Christie K, Strauss SY. 2019. Reproductive isolation and the maintenance of species boundaries in two serpentine endemic Jewelflowers. Evolution, 73: 1375–1391.

Coyne JA, Orr HA. 2004. Speciation. Sunderland, MA: Sinauer Associates.

Cramer MD, West AG, Power SC, Skelton R, Stock WD. 2014. Plant ecophysiological diversity. In: Allsopp N, Colville JF, Verboom GA, eds. Fynbos: ecology, evolution and conservation of a megadiverse region. Oxford: Oxford University Press.

Cramer MD, Wootton LM, Mazijk R, Verboom GA. 2019. New regionally modelled soil layers improve prediction of vegetation type relative to that based on global soil models. Diversity and Distributions, 25: 1736–1750.

da Silva AR, Malafaia G, Menezes IPP. 2017. Biotools: an R function to predict spatial gene diversity via an individual-based approach. Genetics and Molecular Research 16: 1–6.

de Camargo MGG, Lunau K, Batalha MA, Brings S, de Brito VLG, Morellato LPC. 2019. How flower colour signals allure bees and hummingbirds: a community-level test of the bee avoidance hypothesis. New Phytologist, 222: 1112–1122.

Dellinger AS, Lagomarsino L, Michelangeli F, Dullinger S, Smith SD. 2024. The sequential direct and indirect effects of mountain uplift, climatic niche, and floral trait evolution on diversification dynamics in an Andean plant clade. Systematic Biology, 73: 594–612.

Delpierre GR, du Plessis NM. 1974. The winter-growing Gladioli of South Africa. London: Nasionale Bokhandel Ltd.

Demarche ML, Miller TJ, Kay KM. 2015. An ultraviolet floral polymorphism associated with life history drives pollinator discrimination in Mimulus guttatus. American Journal of Botany, 102: 396–406.

Dormann CF, B. G, Fruend J. 2008. Introducing the bipartite package: analysing ecological networks. R news, 8.

Dyer AG. 2006. Discrimination of flower colours in natural settings by bumblebee species Bombus terrestris (Hymenoptera: Apidea). Entomologia Generalis, 28: 257–268.

Elith J, Phillips SJ, Hastie T, Dudík M, Chee YE, Yates CJ. 2010. A statistical explanation of MaxEnt for ecologists. Diversity and Distributions, 17: 43–57.

Ellis AG, Verboom GA, van der Niet T, Johnson SD, Linder HP. 2014. Speciation and extinction in the greater Cape Floristic Region. In: Allsopp N, Colville JF, Verboom GA, eds. Fynbos: ecology, evolution and conservation of a megadiverse region. Oxford: Oxford University Press.

Elzinga JA, Atlan A, Biere A, Gigord L, Weis AE, Bernasconi G. 2007. Time after time: flowering phenology and biotic interactions. Trends in Ecology & Evolution, 22: 432–439.

Farnitano MC, Sweigart AL. 2023. Strong postmating reproductive isolation in Mimulus section Eunanus. Journal of Evolutionary Biology, 36: 1393–1410.

Fernandez-Mazuecos M, Glover BJ. 2024. Climatic and edaphic niche shifts during plant radiation in the Mediterranean biodiversity hotspot. Annals of Botany.

Finnell LM, Koski MH. 2021. A test of Sensory Drive in plant-pollinator interactions: heterogeneity in the signalling environment shapes pollinator preference for a floral visual signal. New Phytologist, 232: 1436–1448.

Forest F, Goldblatt P, Manning JC, et al. 2014. Pollinator shifts as triggers of speciation in painted petal irises (Lapeirousia: Iridaceae). Annals of Botany, 113: 357–371.

Fox J, Weisberg S. 2019. An {R} Companion to applied regression. Thousand Oaks CA: Sage.

Garcia JE, Hannah L, Shrestha M, Burd M, Dyer AG. 2022. Fly pollination drives convergence of flower coloration. New Phytologist, 233: 52–61.

Goldblatt P, Manning J. 1998. Gladiolus in Southern Africa. Vlaeberg: Fernwood Press.

Goldblatt P, Manning JC, Bernhardt P. 2001. Radiation of pollination systems in Gladiolus (Iridaceae: Crocoideae) in southern Africa. Annals of the Missouri Botanical Garden, 88: 713–734.

Grossenbacher DL, Veloz SD, Sexton JP. 2014. Niche and range size patterns suggest that speciation begins in small, ecologically diverged populations in North American monkeyflowers (Mimulus spp.). Evolution, 68: 1270–1280.

Hannah L, Dyer AG, Garcia JE, Dorin A, Burd M. 2019. Psychophysics of the hoverfly: categorical or continuous color discrimination? Current Zoology, 65: 483–492.

Herve M. 2023. RVAideMemoire: testing and plotting procedures for biostatistics. R package version 0.9-83-7. https://CRAN.R-project.org/package=RVAideMemoire.

Hijmans RJ. 2024. raster: geographic data analysis and modeling. R package version 3.6-30. https://rspatial.org/raster.

Hijmans RJ, Phillips S, Leathwick J. 2023. dismo: species distribution modeling. R package version 1.3-15. https://github.com/rspatial/dismo.

Ivey CT, Habecker NM, Bergmann JP, Ewald J, Frayer ME, Coughlan JM. 2023. Weak reproductive isolation and extensive gene flow between Mimulus glaucescens and M. guttatus in northern California. Evolution, 77: 1245–1261.

Johnson SD. 2010. The pollination niche and its role in the diversification and maintenance of the southern African flora. Philosophical Transactions of the Royal Society B, 365: 499–516.

Kay KM. 2006. Reproductive isolation between two closely related hummingbird-pollinated neotropical gingers. Evolution, 60: 538–552.

Latimer AM, Silander Jr. JA, Rebelo AG, Midgley GF. 2009. Experimental biogeography: the role of environmental gradients in high geographic diversity in Cape Proteaceae. Oecologia, 160: 151–162.

Lenth RV. 2022. emmeans: estimated marginal means, aka least-squares means. R package version 1.7.2. https://CRAN.R-project.org/package=emmeans.

Lewis GJ, Obermeyer AA, Barnard TT. 1972. Gladiolus a revision of the South African species. Cape Town: Purnell & Sons.

Linder HP. 2003. The radiation of the Cape flora, southern Africa. Biological Reviews, 78: 597–638.

Linder HP. 2020. The evolution of flowering phenology: an example from the wind-pollinated African Restionaceae. Annals of Botany, 126: 1141–1153.

Lowry DB. 2012. Ecotypes and the controversy of stages in the formation of new species. Biological Journal of the Linnean Society, 106: 241–257.

Maia R, Gruson H, Endler JA, White TE, O’Hara RB. 2019. pavo 2: new tools for the spectral and spatial analysis of colour in R. Methods in Ecology and Evolution, 10: 1097–1107.

Maia R, White TE. 2018. Comparing colors using visual models. Behavioral Ecology, 29: 649–659.

Manning J, Goldblatt P. 2012. Plants of the Greater Cape Floristic Region 1: the core Cape flora. Pretoria: Strelitzia 29.

Minnaar C, de Jager ML, Anderson B. 2019. Intraspecific divergence in floral-tube length promotes asymmetric pollen movement and reproductive isolation. New Phytologist, 224: 1160–1170.

Mitchell N, Moore TE, Mollmann HK, et al. 2015. Functional traits in parallel evolutionary radiations and trait-environment associations in the Cape Floristic Region of South Africa. The American Naturalist, 185: 525–537.

Moir M, Butler H, Peter C, Dold T, Newman E. 2025. A test of the Grant-Stebbins pollinator-shift model of floral evolution. New Phytologist, 245: 2322–2335.

Munguia-Rosas MA, Ollerton J, Parra-Tabla V, De-Nova JA. 2011. Meta-analysis of phenotypic selection on flowering phenology suggests that early flowering plants are favoured. Ecology Letters, 14: 511–521.

Newman E, Johnson SD. 2021. A shift in long-proboscid fly pollinators and floral tube length among populations of Erica junonia (Ericaceae). South African Journal of Botany, 142: 451–458.

Newman E, Johnson SD. 2024. Pollinator-mediated isolation promotes coexistence of closely related food-deceptive orchids. Journal of Evolutionary Biology, 38: 190–201.

Newman E, Manning J, Anderson B. 2014. Matching floral and pollinator traits through guild convergence and pollinator ecotype formation. Annals of Botany, 113: 373–384.

Newman E, Manning J, Anderson B. 2015. Local adaptation: mechanical fit between floral ecotypes of Nerine humilis (Amaryllidaceae) and pollinator communities. Evolution, 69: 2262–2275.

Newman EL, Khoury KL, Niekerk SEV, Peter CI. 2022. Structural anther mimics improve reproductive success through dishonest signaling that enhances both attraction and the morphological fit of pollinators with flowers. Evolution, 76: 1749–1761.

Oksanen J, Blanchet FG, Friendly M, et al. 2020. vegan: community ecology package. R package version 2.5-7. https://CRAN.R-project.org/package=vegan.

Ostevik KL, Andrew RL, Otto SP, Rieseberg LH. 2016. Multiple reproductive barriers separate recently diverged sunflower ecotypes. Evolution, 70: 2322–2335.

Paudel BR, Burd M, Shrestha M, Dyer AG, Li QJ. 2018. Reproductive isolation in alpine gingers: how do coexisting Roscoea (R. purpurea and R. tumjensis) conserve species integrity? Evolution, 72: 1840–1850.

Pewsey A, Neuhäuser M, G.D. R. 2013. Circular statistics in R. Oxford: Oxford University Press.

Phillips SJ, Anderson RP, Schapire RE. 2006. Maximum entropy modeling of species geographic distributions. Ecological Modelling, 190: 231–259.

R Core Team. 2021. R: A language and environment for statistical computing. Vienna, Austria: R Foundation for Statistical Computing.

Ramirez-Aguirre E, Marten-Rodriguez S, Quesada-Avila G, et al. 2019. Reproductive isolation among three sympatric Achimenes species: pre- and post-pollination components. American Journal of Botany, 106: 1021–1031.

Ramsey J, Bradshaw HD, Schemske DW. 2003. Components of reproductive isolation between the monkeyflowers Mimulus lewisii and M. cardinalis (Phrymaceae). Evolution, 57: 1520–1534.

Rundle HD, Nosil P. 2005. Ecological speciation. Ecology Letters, 8: 336–352.

Sandstedt GD, Wu CA, Sweigart AL. 2021. Evolution of multiple postzygotic barriers between species of the Mimulus tilingii complex. Evolution, 75: 600–613.

Schluter D. 2000. The ecology of adaptive radiation. New York: OUP Oxford.

Schluter D. 2001. Ecology and the origin of species. Trends in Ecology & Evolution, 16: 372–380.

Schnitzler J, Barraclough TG, Boatwright JS, et al. 2011. Causes of plant diversification in the Cape biodiversity hotspot of South Africa. Systematic Biology, 60: 343–357.

Sobel JM. 2014. Ecogeographic isolation and speciation in the genus Mimulus. The American Naturalist, 184: 565–579.

Sobel JM, Chen GF. 2014. Unification of methods for estimating the strength of reproductive isolation. Evolution, 68: 1511–1522.

Sobel JM, Streisfeld MA. 2015. Strong premating reproductive isolation drives incipient speciation in Mimulus aurantiacus. Evolution, 69: 447–461.

Troje N. 1993. Spectral categories in the learning behaviour of blowflies. Zeitschrift für Naturforschung C, 48: 96–104.

Valente LM, Manning JC, Goldblatt P, Vargas P. 2012. Did pollination shifts drive diversification in southern African Gladiolus? Evaluating the model of pollinator-driven speciation. The American Naturalist, 180: 83–98.

van der Niet T, Johnson SD. 2009. Patterns of plant speciation in the Cape Floristic Region. Molecular Phylogenetics and Evolution, 51: 85–93.

