## Supplementary materials for "Ecological niche differentiation mediates near complete premating reproductive isolation within the *Gladiolus carneus* (Iridaceae) species complex"

**Table S1**. Coordinates, and elevation (m) of all 29 *G. carneus* sites sampled for this manuscript. The codes for each site are used throughout the manuscript.

| Variety | Site | Code | Latitude | Longitude | Elevation (m) |
|---|---|---|---|---|---|
| *albidus* | Bainskloof A | BK | -33.641822 | 19.075238 | 337 |
| *albidus* | Bainskloof B | BL | -33.639557 | 19.080812 | 372 |
| *albidus* | Franschhoek Pass | FP | -33.921289 | 19.141642 | 510 |
| *albidus* | Helderberg | HB | -34.054097 | 18.874192 | 216 |
| *albidus* | Hermanus | HM | -34.416530 | 19.227500 | 48 |
| *albidus* | Limietberg | LB | -33.707683 | 19.053684 | 457 |
| *albidus* | Stellenboschberg | SB | -33.943373 | 18.886395 | 210 |
| *albidus* | Shaw's Pass | SP | -34.305006 | 19.417495 | 339 |
| *blandus* | Kleinmond Coast | KC | -34.345452 | 19.008462 | 9 |
| *callistus* | Brodie-link C | BC | -34.361673 | 18.865795 | 17 |
| *callistus* | Kleinmond Golf Course | KG | -34.326530 | 19.030778 | 74 |
| *callistus* | Pringle Bay C | PB | -34.343890 | 18.847427 | 19 |
| *high-altitude* | Jonkershoek | JH | -34.001640 | 19.012690 | 1235 |
| *high-altitude* | Jonaskop | JK | -33.963432 | 19.510247 | 1403 |
| *high-altitude* | Mont Rochelle | MR | -33.882298 | 19.173532 | 1303 |
| *langeberg* | Garcia's Pass | GP | -33.957285 | 21.201655 | 749 |
| *langeberg* | Prince Alfred's Pass | PA | -33.801141 | 23.178886 | 366 |
| *langeberg* | Tradouw Pass | TP | -33.950402 | 20.702005 | 249 |
| *macowanianus* | Brodie-Link M/Betty's Bay | BM | -34.362433 | 18.866515 | 16 |
| *macowanianus* | Devil's Peak | DP | -33.956420 | 18.425135 | 475 |
| *macowanianus* | Elsie's Peak | EP | -34.146345 | 18.430717 | 211 |
| *macowanianus* | Franskraal/Gansbaai | FK | -34.600648 | 19.394002 | 17 |
| *macowanianus* | Flower Valley Conservation | FV | -34.548015 | 19.472299 | 122 |
| *macowanianus* | Lomond Wine Farm | LW | -34.567372 | 19.481217 | 27 |
| *macowanianus* | Pringle Bay M | PM | -34.348830 | 18.832518 | 18 |
| *macowanianus* | Rhodes Memorial | RM | -33.950491 | 18.455636 | 231 |
| *prismatosiphon* | Bredasdorp | BD | -34.544306 | 20.035664 | 225 |
| *prismatosiphon* | Grootbos/Farm 215 | GB | -34.553089 | 19.546951 | 262 |
| *prismatosiphon* | Napier | NP | -34.509889 | 19.879167 | 404 |

**Table S2.** Sample sizes of the morphological trait measurements at each *G. carneus* study population. Varieties include those based on former descriptions by Delpierre and du Plessis (1974) and the two newly described varieties of the species.

| Population number | Variety | Site | tube length (mm) | flower gape (mm) | petal size (mm) | flower width (mm) | inflore scence height (cm) | total flowers | total leaves | leaf length (cm) | leaf width (cm) |
|---|---|---|---|---|---|---|---|---|---|---|---|
| 1 | *albidus* | Bainskloof A | 5 | 5 | 0 | 5 | 0 | 5 | 0 | 0 | 0 |
| 2 | *albidus* | Bainskloof B | 7 | 6 | 0 | 7 | 0 | 7 | 0 | 0 | 0 |
| 3 | *albidus* | Franschhoek pass | 9 | 9 | 0 | 8 | 0 | 8 | 0 | 0 | 0 |
| 4 | *albidus* | Helderberg | 35 | 31 | 26 | 29 | 20 | 28 | 20 | 20 | 20 |
| 5 | *albidus* | Hermanus | 5 | 4 | 5 | 3 | 5 | 5 | 5 | 2 | 5 |
| 6 | *albidus* | Limietberg | 34 | 30 | 17 | 30 | 16 | 30 | 16 | 15 | 15 |
| 7 | *albidus* | Shaw's Pass | 8 | 8 | 8 | 8 | 0 | 10 | 0 | 0 | 0 |
| 8 | *albidus* | Stellenboschberg | 11 | 8 | 11 | 11 | 11 | 11 | 11 | 9 | 10 |
| 9 | *blandus* | Kleinmond Coast | 38 | 37 | 21 | 38 | 22 | 31 | 22 | 22 | 22 |
| 10 | *callistus* | Brodie-Link C | 26 | 24 | 26 | 26 | 20 | 20 | 20 | 20 | 20 |
| 11 | *callistus* | Kleinmond Golf Course | 32 | 31 | 32 | 29 | 23 | 23 | 23 | 20 | 22 |
| 12 | *callistus* | Pringle Bay C | 9 | 10 | 0 | 10 | 0 | 10 | 0 | 0 | 0 |
| 13 | *high-altitude* | Jonaskop | 31 | 31 | 31 | 30 | 15 | 30 | 15 | 15 | 15 |
| 14 | *high-altitude* | Jonkershoek | 16 | 17 | 17 | 14 | 7 | 17 | 7 | 5 | 7 |
| 15 | *high-altitude* | Mont Rochelle | 40 | 35 | 38 | 29 | 32 | 40 | 15 | 15 | 15 |
| 16 | *langeberg* | Garcias pass | 16 | 14 | 0 | 17 | 0 | 17 | 0 | 0 | 0 |
| 17 | *langeberg* | Tradouw pass | 40 | 39 | 29 | 40 | 10 | 40 | 10 | 10 | 10 |
| 18 | *macowanianus* | Betty's Bay | 27 | 27 | 10 | 27 | 10 | 27 | 10 | 10 | 10 |
| 19 | *macowanianus* | Brodie-Link M | 15 | 12 | 15 | 12 | 10 | 10 | 10 | 10 | 9 |
| 20 | *macowanianus* | Devil's Peak | 28 | 25 | 25 | 25 | 20 | 25 | 20 | 20 | 19 |
| 21 | *macowanianus* | Elsie's Peak | 35 | 34 | 24 | 33 | 20 | 30 | 20 | 20 | 20 |
| 22 | *macowanianus* | Flower Valley Conservation Trust | 26 | 26 | 26 | 0 | 0 | 0 | 0 | 0 | 0 |

| | | | | | | | | | | | |
|---|---|---|---|---|---|---|---|---|---|---|---|
| 23 | *macowanianus* | Franskraal | 34 | 30 | 18 | 31 | 14 | 30 | 14 | 8 | 13 |
| 24 | *macowanianus* | Lomond Wine Farm | 8 | 8 | 8 | 7 | 0 | 8 | 0 | 0 | 0 |
| 25 | *macowanianus* | Rhodes Memorial | 35 | 34 | 34 | 34 | 20 | 20 | 20 | 20 | 20 |
| 26 | *prismatosiphon* | Bredasdorp | 21 | 19 | 20 | 18 | 15 | 15 | 15 | 15 | 15 |
| 27 | *prismatosiphon* | Grootbos | 26 | 26 | 26 | 25 | 20 | 20 | 20 | 20 | 19 |
| 28 | *prismatosiphon* | Napier | 23 | 23 | 23 | 22 | 0 | 0 | 0 | 0 | 0 |
| **Total** | | | **640** | **603** | **490** | **568** | **310** | **517** | **293** | **276** | **286** |

**Table S3**. Sample sizes of individuals for which spectral reflectance were collected for each *G. carneus* study population and variety.

| Site number | Variety | Site | Number of individuals | Year collected |
|---|---|---|---|---|
| 1 | *albidus* | Bainskloof A | 5 | 2020 |
| 2 | *albidus* | Franschhoek Pass | 7 | 2020 |
| 3 | *albidus* | Helderberg | 18 | 2020, 2023 |
| 4 | *albidus* | Hermanus | 3 | 2023 |
| 5 | *albidus* | Limietberg | 14 | 2020 |
| 6 | *albidus* | Stellenboschberg | 6 | 2023 |
| 7 | *blandus* | Kleinmond Coast | 22 | 2020, 2023 |
| 8 | *callistus* | Brodie-link C | 12 | 2023 |
| 9 | *callistus* | Kleinmond golf course | 14 | 2023 |
| 10 | *callistus* | Pringle Bay C | 6 | 2020 |
| 11 | *high-altitude* | Jonaskop | 14 | 2022, 2023 |
| 12 | *high-altitude* | Jonkershoek | 10 | 2023 |
| 13 | *high-altitude* | Mont Rochelle | 8 | 2023 |
| 14 | *langeberg* | Garcia's Pass | 16 | 2020 |
| 15 | *langeberg* | Tradouw Pass | 10 | 2020 |
| 16 | *macowanianus* | Betty's Bay | 28 | 2020, 2023 |
| 17 | *macowanianus* | Devil's Peak | 11 | 2023 |
| 18 | *macowanianus* | Elsie's Peak | 19 | 2020, 2023 |
| 19 | *macowanianus* | Franskraal/Gansbaai | 16 | 2020 |
| 20 | *macowanianus* | Pringle bay M | 10 | 2020 |
| 21 | *macowanianus* | Rhodes Memorial | 12 | 2023 |
| 22 | *prismatosiphon* | Bredasdorp | 12 | 2023 |
| 23 | *prismatosiphon* | Grootbos | 12 | 2023 |
| 24 | *prismatosiphon* | Napier | 15 | 2022 |
| ***Subtotals*** | | | | |
| | *albidus* | | 53 | |
| | *blandus* | | 22 | |
| | *callistus* | | 32 | |
| | *high-altitude* | | 32 | |
| | *langeberg* | | 26 | |
| | *macowanianus* | | 96 | |
| | *prismatosiphon* | | 39 | |
| **Total** | | | 300 | |

**Table S4**. Bioclimatic and topography layers mined from Worldclim and the soil layers from Cramer *et al*. (2019) that were used in the abiotic niche analysis. Bioclimatic, soil, and elevational layers in bold were the uncorrelated variables used in the niche modelling and differentiation analysis.

| Layer category | Abiotic layers | Description | Unit |
|---|---|---|---|
| Bioclimatic | Bio1 | Annual Mean Temperature | °C |
| | Bio2 | Mean Diurnal Range (Mean of monthly (max temp - min temp)) | °C |
| | **Bio3** | **Isothermality (BIO2/BIO7) (×100)** | % |
| | Bio4 | Temperature Seasonality (standard deviation ×100) | - |
| | Bio5 | Max Temperature of Warmest Month | °C |
| | Bio6 | Min Temperature of Coldest Month | °C |
| | **Bio7** | **Temperature Annual Range (BIO5-BIO6)** | °C |
| | Bio8 | Mean Temperature of Wettest Quarter | °C |
| | Bio9 | Mean Temperature of Driest Quarter | °C |
| | Bio10 | Mean Temperature of Warmest Quarter | °C |
| | Bio11 | Mean Temperature of Coldest Quarter | °C |
| | **Bio12** | **Annual Precipitation** | mm |
| | Bio13 | Precipitation of Wettest Month | mm |
| | Bio14 | Precipitation of Driest Month | mm |
| | **Bio15** | **Precipitation Seasonality (Coefficient of Variation)** | - |
| | Bio16 | Precipitation of Wettest Quarter | mm |
| | Bio17 | Precipitation of Driest Quarter | mm |
| | Bio18 | Precipitation of Warmest Quarter | mm |
| | Bio19 | Precipitation of Coldest Quarter | mm |
| Soil | pH | pH | - |
| | **EC** | **electrical conductivity** | mS/m |
| | total C | total Carbon | %, w/w |
| | **total N** | **total Nitrogen** | %, w/w |
| | **extractable P** | **extractable Phosphorus** | mg/kg |
| | extractable K | extractable Potassium | $cmol^{+}kg^{-1}$ |
| | **extractable NA** | **extractable Sodium** | $cmol^{+}kg^{-1}$ |
| Topography | | **Elevation** | meters above sea level |

**Table S5**. Mean and sample size of visitation rates and pollen loads for each functional pollinator collected across populations and varieties of *G. carneus*.

| Variety | Site | Site Code | Functional pollinator | Visitation rate (visits.flower$^{-1}$.hour$^{-1}$) | | | | Pollen loads | Pollinators caught |
|---|---|---|---|---|---|---|---|---|---|
| | | | | Mean | Sample size | Obs flowers | Obs time (hours) | Mean | Total |
| *albidus* | Bainskloof A | BK | Solitary bees | 0.133 | 1 | 5 | 3.00 | - | - |
| *albidus* | Helderberg | HB | Solitary bees | 0.065 | 2 | 51 | 4.98 | 38 | 2 |
| *albidus* | Limietberg | LB | Carpenter bees | 0.035 | 1 | 52 | 1.10 | - | - |
| *albidus* | Limietberg | LB | Solitary bees | 0.142 | 3 | 190 | 10.45 | 104 | 4 |
| *callistus* | Brodie-Link C | BC | Carpenter bees | 0.014 | 1 | 66 | 3.27 | - | - |
| *callistus* | Brodie-Link C | BC | Honey bees | 0.009 | 1 | 66 | 3.27 | - | - |
| *callistus* | Brodie-Link C | BC | MTFs | 0.283 | 1 | 66 | 3.27 | 1549 | 5 |
| *callistus* | Brodie-Link C | BC | Solitary bees | 0.079 | 1 | 66 | 3.27 | 36 | 4 |
| *callistus* | Kleinmond Golf Course | KG | LTFs | 0.006 | 1 | 42 | 4.32 | 139 | 1 |
| *callistus* | Kleinmond Golf Course | KG | Solitary bees | 0.065 | 2 | 50 | 7.97 | 62 | 5 |
| *callistus* | Pringle Bay | PB | Solitary bees | 0.571 | 1 | 14 | 1.00 | - | - |
| *high-altitude* | Jonaskop | JK | LTFs | 0.202 | 1 | 23 | 3.23 | 1175 | 8 |
| *high-altitude* | Jonaskop | JK | MTFs | - | - | - | - | 1117 | 1 |
| *high-altitude* | Jonaskop | JK | Solitary bees | 0.121 | 1 | 23 | 3.23 | 68 | 2 |
| *high-altitude* | Mont Rochelle | MR | Solitary bees | 1.000 | 1 | 7 | 1.00 | 1065 | 1 |
| *langeberg* | Tradouw Pass | TP | Honey bees | 0.344 | 1 | 30 | 3.00 | - | - |
| *langeberg* | Tradouw Pass | TP | Lycaenid butterfly | 0.011 | 1 | 30 | 3.00 | - | - |
| *langeberg* | Tradouw Pass | TP | Solitary bees | 0.084 | 2 | 46 | 5.00 | - | - |
| *macowanianus* | Brodie-Link M | BM | Honey bees | - | - | - | - | 14 | 1 |
| *macowanianus* | Brodie-Link M | BM | MTFs | - | - | - | - | 731 | 7 |
| *macowanianus* | Devils Peak | DP | MTFs | 0.170 | 3 | 45 | 7.72 | - | - |
| *macowanianus* | Elsies Peak | EP | LTFs | 0.214 | 1 | 7 | 2.00 | 16 | 1 |

| | | | | | | | | | |
|---|---|---|---|---|---|---|---|---|---|
| *macowanianus* | Elsies Peak | EP | MTFs | - | - | - | - | 162 | 1 |
| *macowanianus* | Elsies Peak | EP | Solitary bees | 0.074 | 1 | 21 | 3.22 | - | - |
| *macowanianus* | Rhodes Memorial | RM | Solitary bees | 0.049 | 2 | 32 | 8.28 | 35 | 2 |
| *prismatosiphon* | Bredasdorp | BD | Solitary bees | 0.031 | 1 | 156 | 3.08 | 54 | 2 |
| *prismatosiphon* | Grootbos | GB | LTFs | 0.063 | 2 | 23 | 7.20 | 1197 | 2 |
| *prismatosiphon* | Napier | NP | LTFs | - | - | - | - | 10 | 1 |
| **Total** | | | | | | **729** | **69.40** | | |

**Table S6**. Generalised linear model outputs testing for differences between the *G. carneus* varieties morphological traits. The models included two explanatory variables, variety and site nested within variety.

| | **variety** | | | **variety:site** | | |
|---|---|---|---|---|---|---|
| **Traits** | $\chi^2$ | *df* | *P* | $\chi^2$ | *df* | *P* |
| tube length | 749.82 | 6 | **< 0.0001** | 855.67 | 21 | **< 0.0001** |
| flower gape | 409.09 | 6 | **< 0.0001** | 161.64 | 21 | **< 0.0001** |
| petal size | 118.15 | 6 | **< 0.0001** | 345.85 | 21 | **< 0.0001** |
| inflorescence height | 44.11 | 6 | **< 0.0001** | 125.25 | 12 | **< 0.0001** |
| leaf length | 164.9 | 6 | **< 0.0001** | 222.57 | 12 | **< 0.0001** |
| leaf width | 359.23 | 6 | **< 0.0001** | 294.14 | 12 | **< 0.0001** |

**Table S7.** Pairwise comparisons between the morphological traits of *G. carneus* varieties. Significant differences between pairs of varieties are highlighted in bold.

| Comparisons | Tube length | Flower gape | Petal size | Inflorescence height | Length longest leaf | Width of longest leaf |
|---|---|---|---|---|---|---|
| *albidus - blandus* | **< 0.0001** | **< 0.0001** | 1.000 | 1.000 | 1.000 | 0.8449 |
| *albidus - callistus* | **0.0384** | 1.000 | 1.000 | **0.0004** | **0.0003** | **< 0.0001** |
| *albidus – high-altitude* | 1.000 | 1.000 | 1.000 | **0.0059** | **< 0.0001** | **< 0.0001** |
| *albidus - langeberg* | **< 0.0001** | **< 0.0001** | 0.2624 | **0.0025** | **0.0010** | **0.0001** |
| *albidus - macowanianus* | 0.3563 | **< 0.0001** | **< 0.0001** | 0.0773 | 1.000 | **< 0.0001** |
| *albidus - prismatosiphon* | **< 0.0001** | **0.0002** | **< 0.0001** | 0.0873 | **0.0003** | **0.0032** |
| *blandus - callistus* | **< 0.0001** | **< 0.0001** | 1.000 | **0.0208** | **< 0.0001** | **< 0.0001** |
| *blandus – high-altitude* | **< 0.0001** | **< 0.0001** | 1.000 | 0.1091 | **< 0.0001** | **< 0.0001** |
| *blandus - langeberg* | **< 0.0001** | 0.248 | 1.000 | **0.0056** | **0.0032** | **< 0.0001** |
| *blandus - macowanianus* | **< 0.0001** | **< 0.0001** | 0.4701 | 1.000 | 1.000 | **< 0.0001** |
| *blandus - prismatosiphon* | **< 0.0001** | **< 0.0001** | **0.0003** | 0.6848 | **< 0.0001** | **< 0.0001** |
| *callistus - high-altitude* | **0.0165** | 1.000 | 1.000 | 1.000 | **0.0063** | **0.0002** |
| *callistus - langeberg* | **< 0.0001** | **< 0.0001** | **0.0168** | **< 0.0001** | **< 0.0001** | 1.000 |
| *callistus - macowanianus* | 1.000 | **< 0.0001** | **< 0.0001** | 0.5866 | **< 0.0001** | 0.0774 |
| *callistus - prismatosiphon* | 0.0664 | **0.0009** | **< 0.0001** | 1.000 | 1.000 | **< 0.0001** |
| *high-altitude - langeberg* | **< 0.0001** | **< 0.0001** | 0.3949 | **< 0.0001** | **< 0.0001** | **0.0002** |
| *high-altitude - macowanianus* | 0.1434 | **< 0.0001** | **< 0.0001** | 1.000 | **< 0.0001** | 0.418 |
| *high-altitude - prismatosiphon* | **< 0.0001** | **< 0.0001** | **< 0.0001** | 1.000 | 0.0828 | **< 0.0001** |
| *langeberg - macowanianus* | **< 0.0001** | **0.0055** | 1.000 | **< 0.0001** | **< 0.0001** | **0.0319** |
| *langeberg - prismatosiphon* | **< 0.0001** | **< 0.0001** | **0.009** | **< 0.0001** | **< 0.0001** | 0.1527 |
| *macowanianus - prismatosiphon* | **< 0.0001** | 0.108 | **0.0127** | 1.000 | **< 0.0001** | **< 0.0001** |

**Table S8**. Pairwise comparisons between varieties of *G. carneus* plotted in bee and fly colour vision models. Significant differences are highlighted in bold.

| | Bee colour vision model | | | | | Fly colour vision model | | | | |
|---|---|---|---|---|---|---|---|---|---|---|
| **Comparison** | **Tepal** | **Median tepal centre** | **Median tepal guide** | **Lateral tepal centre** | **Lateral tepal guide** | **Tepal** | **Median tepal centre** | **Median tepal guide** | **Lateral tepal centre** | **Lateral tepal guide** |
| *albidus – blandus* | **< 0.001** | **< 0.001** | - | **< 0.001** | - | **< 0.001** | **< 0.001** | - | **< 0.001** | - |
| *albidus – callistus* | **< 0.001** | - | - | - | - | **< 0.001** | - | - | - | - |
| *albidus – high-altitude* | **< 0.001** | **< 0.001** | - | **< 0.001** | - | **< 0.001** | **< 0.001** | - | **< 0.001** | - |
| *albidus - langeberg* | **< 0.001** | **< 0.001** | - | **< 0.001** | - | **< 0.001** | **< 0.001** | - | **< 0.001** | - |
| *albidus - macowanianus* | **< 0.001** | **< 0.001** | - | **< 0.001** | - | **< 0.001** | **< 0.001** | - | **< 0.001** | - |
| *albidus - prismatosiphon* | **< 0.001** | **< 0.001** | - | **< 0.001** | - | **< 0.001** | **< 0.001** | - | **< 0.001** | - |
| *blandus – callistus* | **< 0.001** | - | - | - | - | **< 0.001** | - | - | - | - |
| *blandus – high-altitude* | **< 0.001** | **0.010** | 1.000 | **< 0.001** | **0.029** | **< 0.001** | **0.004** | 1.000 | **< 0.001** | 0.052 |
| *blandus - langeberg* | **< 0.001** | **< 0.001** | 0.059 | 1.000 | 1.000 | **< 0.001** | **< 0.001** | **0.023** | **0.035** | 1.000 |
| *blandus - macowanianus* | **< 0.001** | **< 0.001** | 1.000 | 0.785 | 0.332 | **0.024** | **< 0.001** | 1.000 | 0.695 | 1.000 |
| *blandus - prismatosiphon* | 0.930 | **< 0.001** | 0.006 | **< 0.001** | 1.000 | 0.793 | **< 0.001** | **0.032** | **< 0.001** | 1.000 |
| *callistus – high-altitude* | 1.000 | - | - | - | - | 1.000 | - | - | - | - |
| *callistus - langeberg* | **< 0.001** | - | - | - | - | **< 0.001** | - | - | - | - |
| *callistus - macowanianus* | **< 0.001** | - | - | - | - | **< 0.001** | - | - | - | - |
| *callistus - prismatosiphon* | **< 0.001** | - | - | - | - | **< 0.001** | - | - | - | - |
| *high-altitude - langeberg* | **< 0.001** | **< 0.001** | 0.028 | **< 0.001** | **< 0.001** | **< 0.001** | **< 0.001** | **0.03** | **< 0.001** | **< 0.001** |
| *high-altitude - macowanianus* | **< 0.001** | 0.251 | 0.872 | **< 0.001** | **0.002** | **< 0.001** | 0.058 | 1.000 | **< 0.001** | **0.001** |
| *high-altitude - prismatosiphon* | **< 0.001** | **< 0.001** | **< 0.001** | **< 0.001** | **< 0.001** | **< 0.001** | **< 0.001** | **0.002** | **< 0.001** | **< 0.001** |
| *langeberg - macowanianus* | **< 0.001** | **< 0.001** | **< 0.001** | 0.051 | 1.000 | **< 0.001** | **< 0.001** | **< 0.001** | **0.002** | 1.000 |
| *langeberg - prismatosiphon* | **< 0.001** | **< 0.001** | 1.000 | 0.002 | 1.000 | **< 0.001** | **< 0.001** | 0.235 | **< 0.001** | 1.000 |
| *macowanianus - prismatosiphon* | **0.002** | **< 0.001** | **< 0.001** | **< 0.001** | 0.627 | **0.018** | **< 0.001** | **< 0.001** | **< 0.001** | 1.000 |

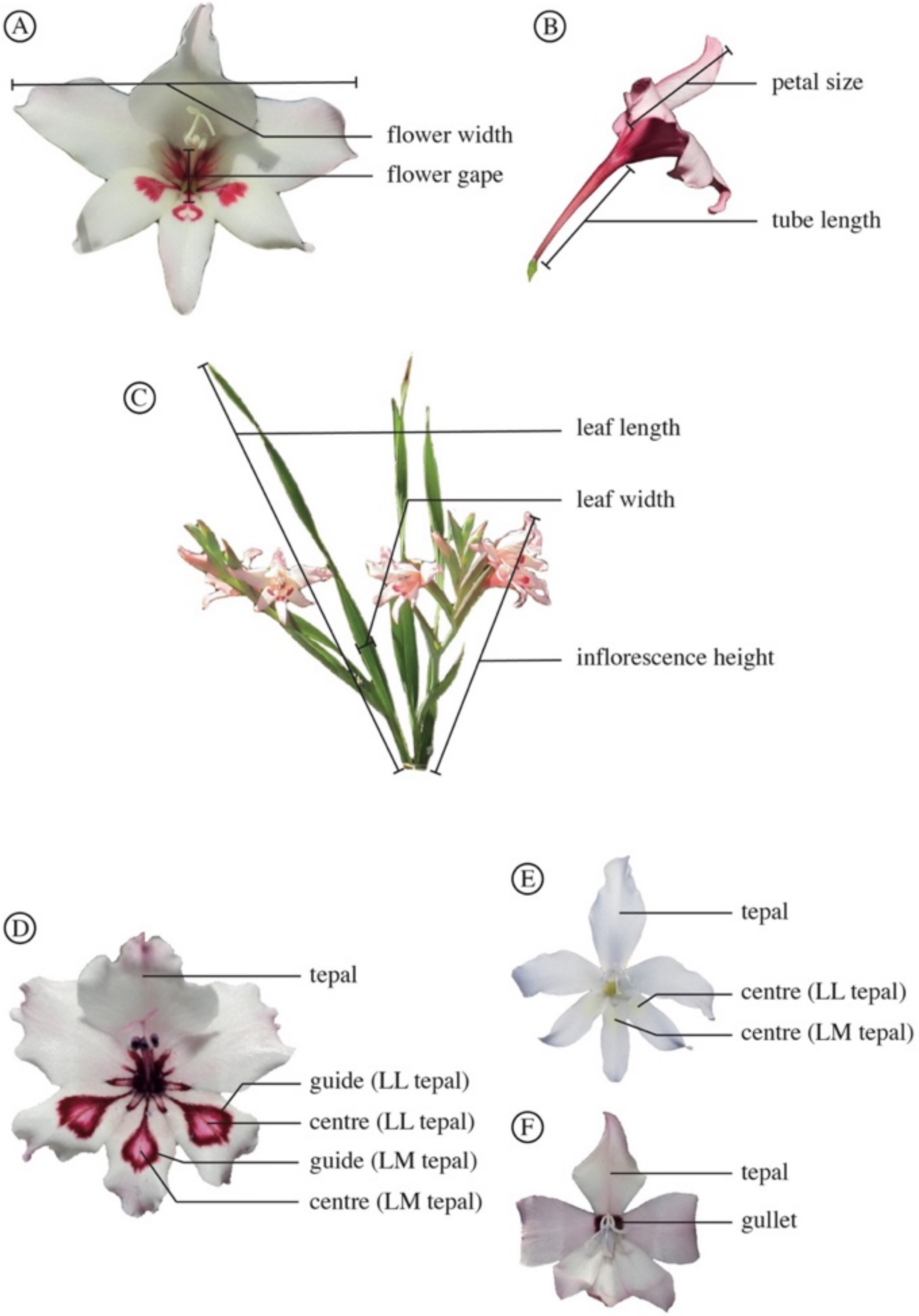


**Figure S1.** Morphological measurements of *G. carneus* varieties. The (A) front view of a *G. carneus* flower showing flower width and flower gape, the (B) side view showing petal size and tube length, (C) whole plant showing measurements of the leaf length, leaf width, and inflorescence height. Additionally, spectral measurements were taken for all varieties from the dorsal tepals, lower median tepal (LM tepal) and lower lateral tepal (LL tepal). Spectral measurements taken for (D) *blandus*, *high-altitude*, *langeberg*, *macowanianus*, *prismatosiphon*, (E) *albidus* and (F) *callistus*.

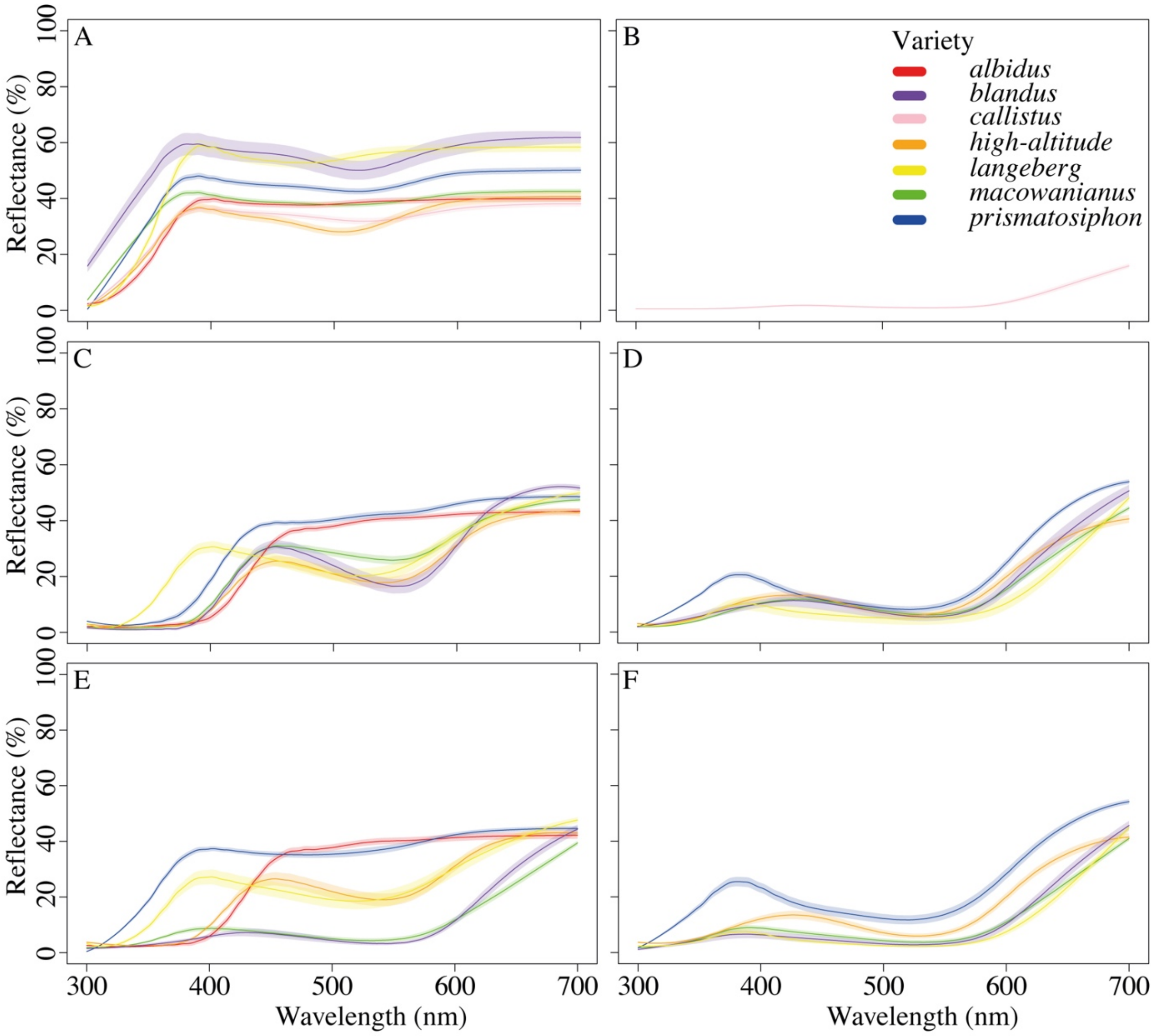


**Figure S2.** Spectral reflectance curves of (A) tepal, (B) gullet, (C) centre of the median tepal, (D) guide of median tepal, (E) centre of lateral tepal, (F) guide of lateral tepal of all *G. carneus* varieties. Mean spectral reflectance curve is depicted by the dark line in the centre with lighter shading representing the standard error.

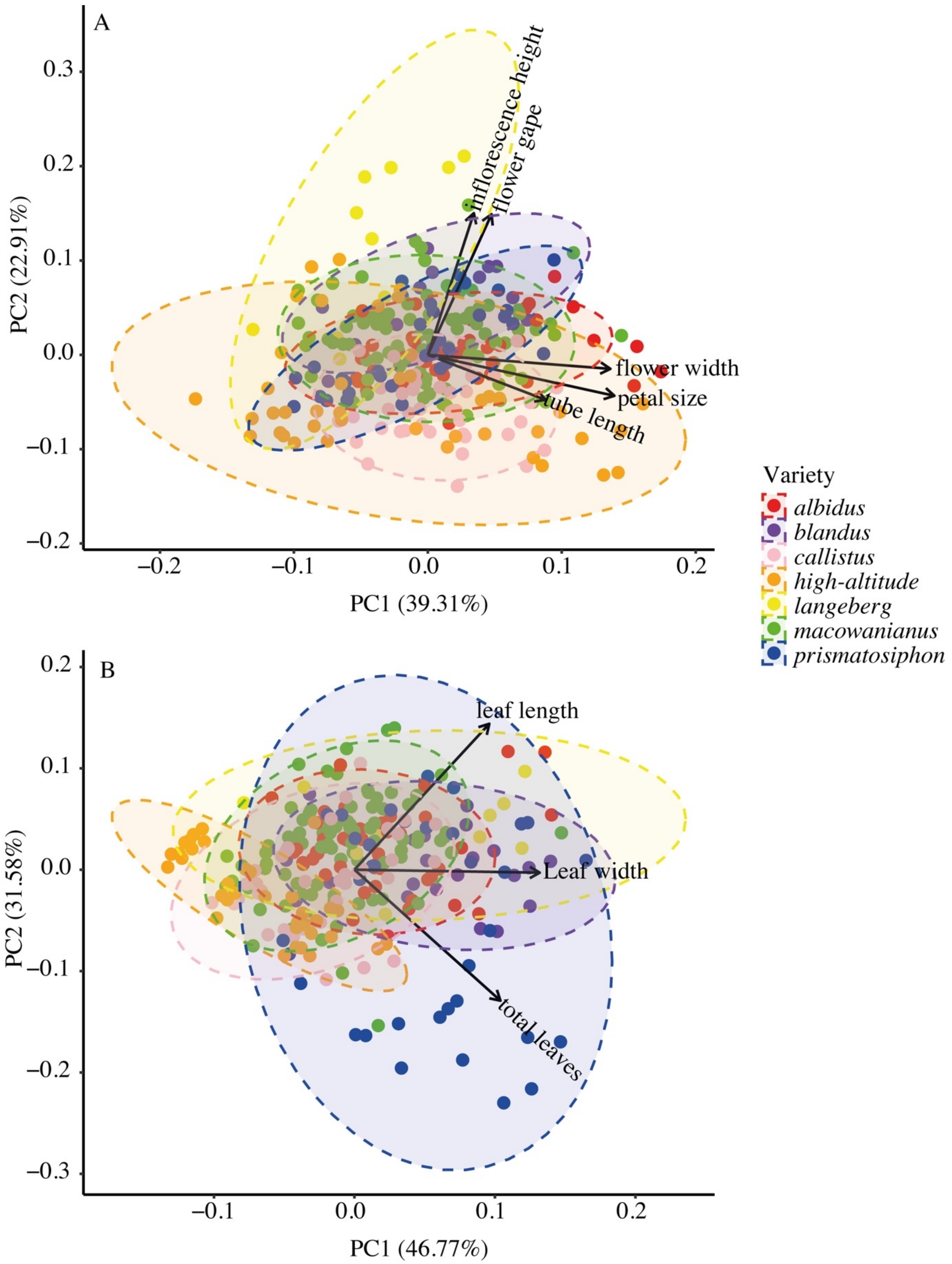


**Figure S3.** PCAs of (A) floral and (B) vegetative traits of the *G. carneus* varieties. The PCA includes ellipses showing 95% confidence intervals and a biplot of trait loadings.

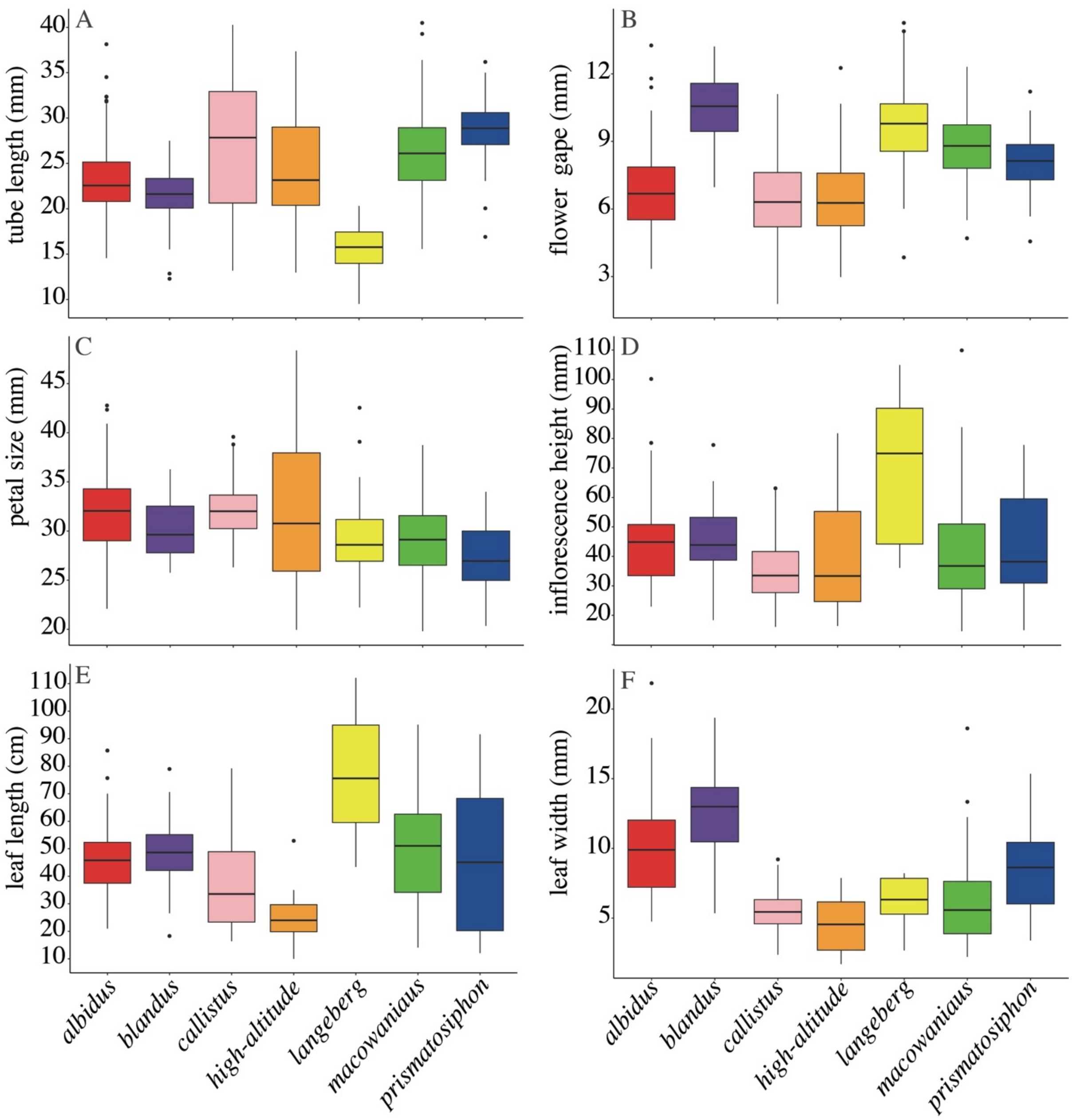


**Figure S4.** Comparisons of the functional traits (A) tube length, (B) flower gape, (C) petal size, (D) inflorescence height, (E) leaf length and (F) leaf width between *G. carneus* varieties.

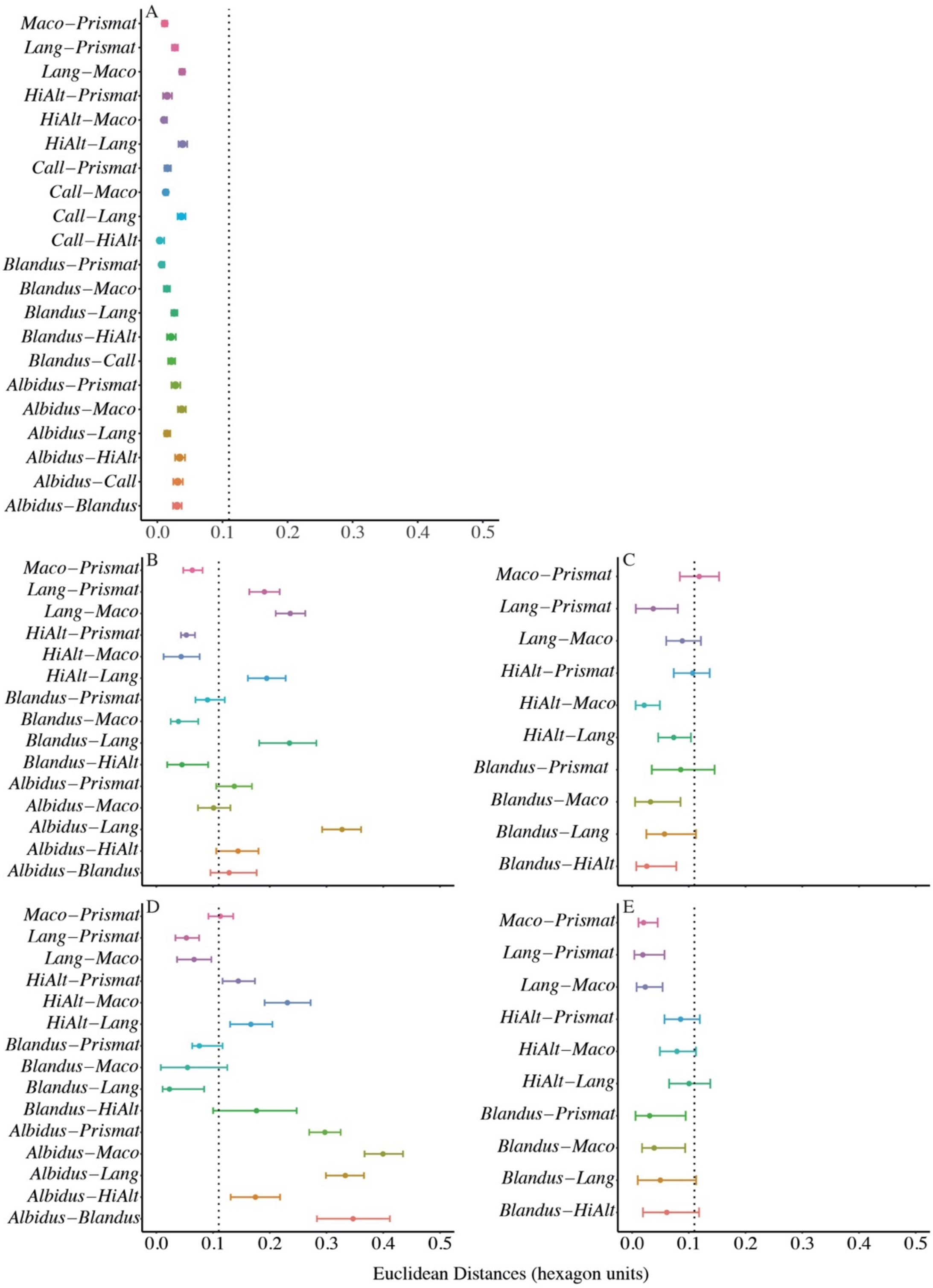


**Figure S5.** Euclidean distances between each variety for the (A) tepal, (B) centre of the median tepal, (C) guide of median tepal, (D) centre of lateral tepal, (E) and guide of lateral tepal modelled in bee colour vision with *Apis mellifera* spectral sensitivities. Dotted lines at 0.11 hexagon units represent the discrimination threshold in bee colour vision. Central point represents the mean with tails representing upper and lower bootstrapped confidence limits.

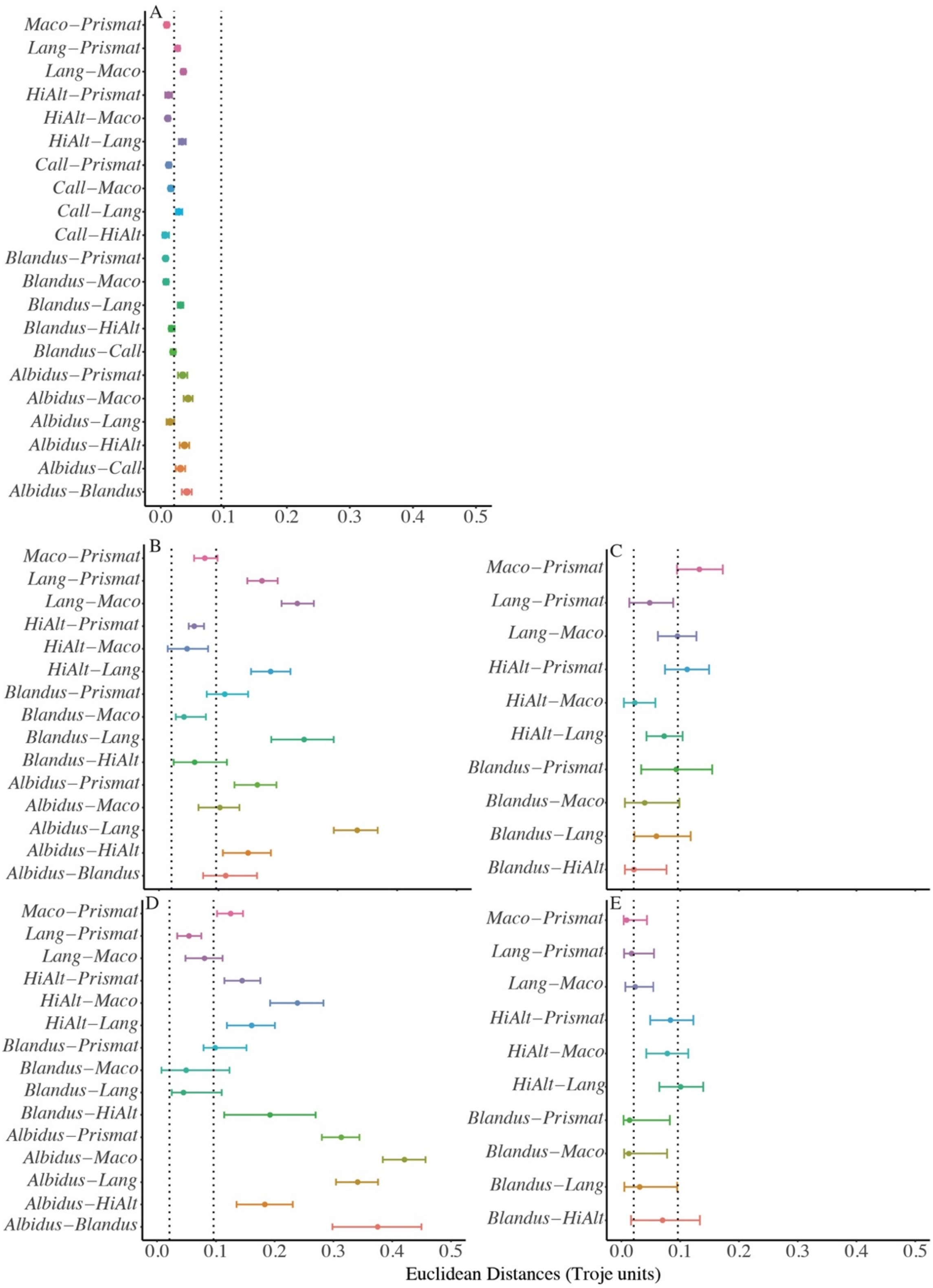


**Figure S6.** Euclidean distances between each variety for (A) tepal, (B) centre of the median tepal, (C) guide of median tepal, (D) centre of lateral tepal, (E) and guide of lateral tepal modelled in fly colour vision with *Eristalis tenax* spectral sensitivities. Dotted lines at 0.021 and 0.096 Troje Units represent the discrimination thresholds in the fly colour vision. Central point represents the mean with tails representing upper and lower bootstrapped confidence limits.

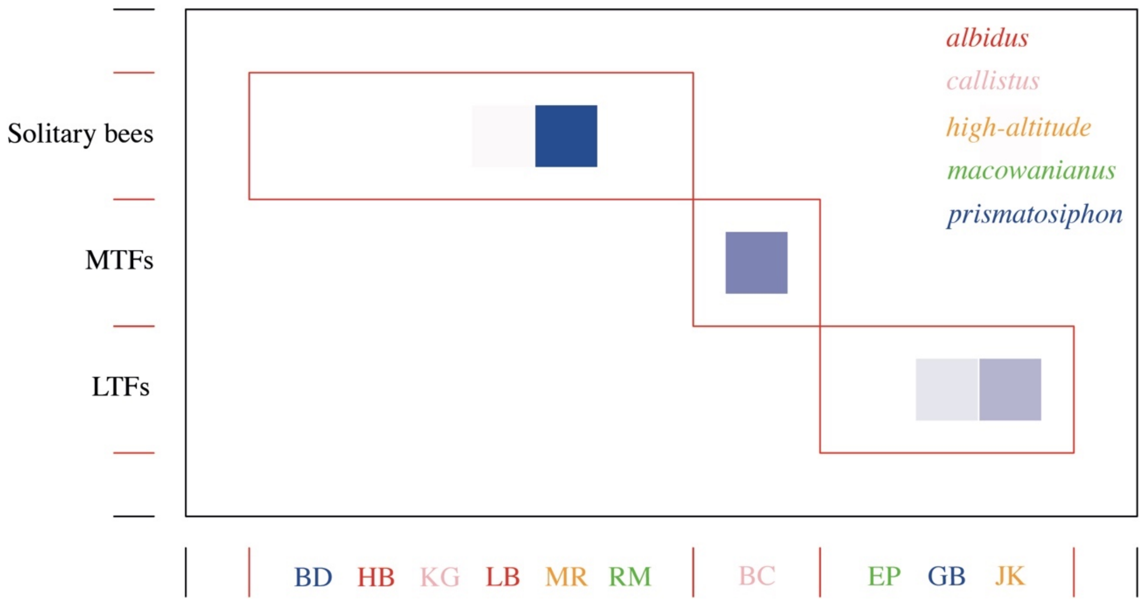


**Figure S7.** Modularity analysis of pollinator importance showing *G. carneus* sites associated with each functional pollinator group.

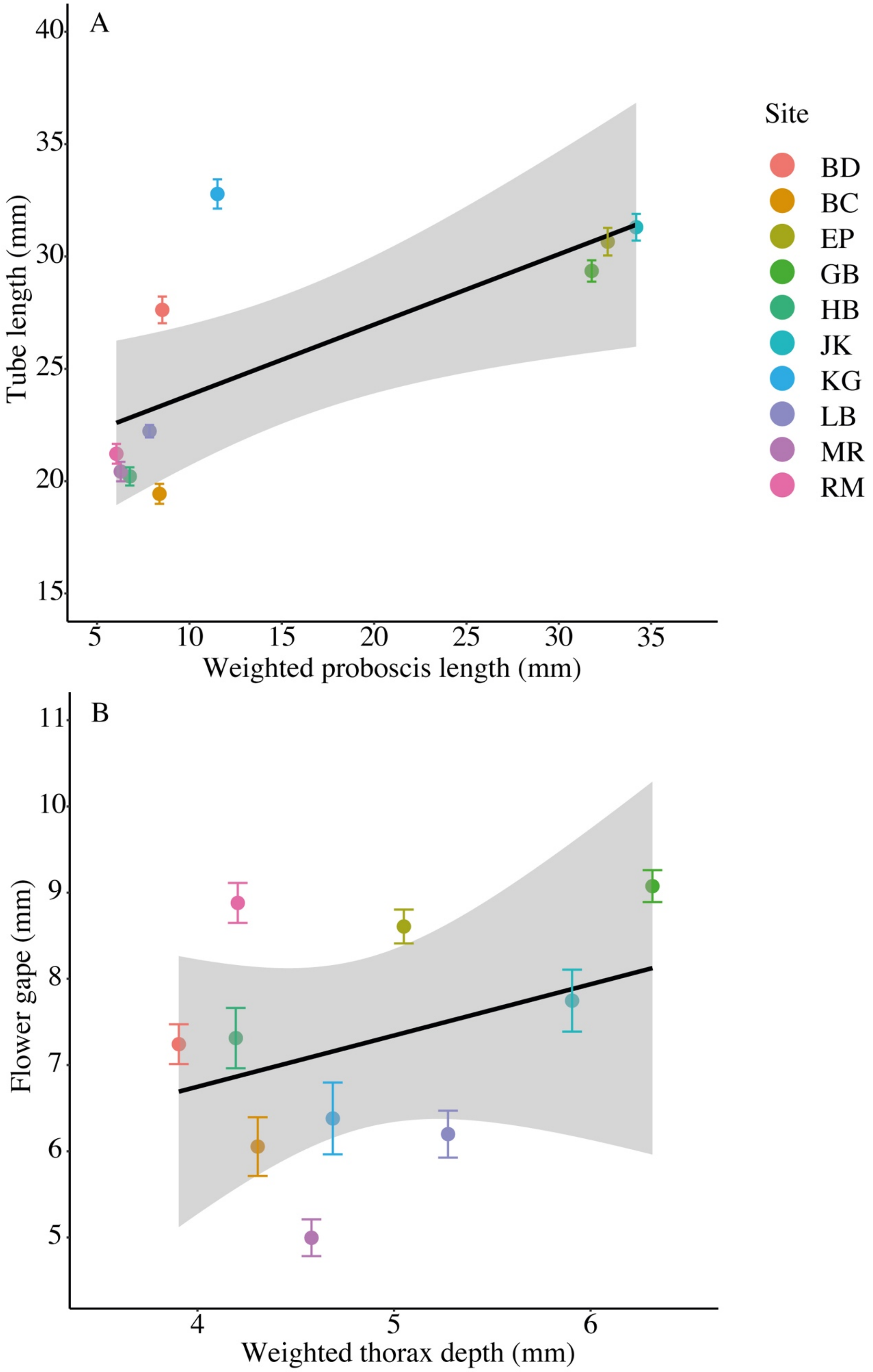


**Figure S8.** Correlations between the (A) weighted proboscis length (mm) of pollinators and *G. carneus* tube length (mean ± SE mm), and (B) weighted thorax depth (mm) of pollinators and *G. carneus* flower gape (mean ± SE mm) across populations of *G. carneus*.
